# Deep learning the cis-regulatory code of chromatin dynamics during cellular reprogramming

**DOI:** 10.1101/2023.10.04.560808

**Authors:** Surag Nair, Mohamed Ameen, Laksshman Sundaram, Anusri Pampari, Jacob Schreiber, Akshay Balsubramani, Yu Xin Wang, David Burns, Helen M Blau, Ioannis Karakikes, Kevin C Wang, Anshul Kundaje

**Affiliations:** Department of Computer Science, Stanford University, Stanford, CA, USA; Department of Cancer Biology, Stanford University, Stanford, CA, USA; Cardiovascular Institute, Stanford University, Stanford, CA, USA; Department of Dermatology, Stanford University, Stanford, CA, USA; Program in Epithelial Biology, Stanford University, Stanford, CA, USA; Department of Genetics, Stanford University, Stanford, CA, USA; Baxter Laboratory for Stem Cell Biology, Stanford University, Stanford, CA, USA; Department of Microbiology and Immunology, Stanford University, Stanford, CA, USA; Department of Cardiothoracic Surgery, Stanford University, Stanford, CA, USA; Veterans Affairs Palo Alto Healthcare System, Palo Alto, CA, USA

## Abstract

The concentration and stoichiometry of transcription factors (TFs) determine cellular identity and can be manipulated to drive cell state transitions. Understanding how changes in TF concentration regulate chromatin state and expression across cell state transitions remains a challenge. We investigated this relationship by profiling chromatin accessibility and gene expression at single-cell resolution across a densely sampled time course of reprogramming human fibroblasts to induced pluripotent stem cells via ectopic expression of *OCT4*, *SOX2*, *KLF4*, and *MYC* (OSKM). Using deep learning sequence models of base-resolution chromatin accessibility profiles across cell states, we deciphered predictive transcription factor (TF) motif syntax in regulatory elements, inferred affinity- and concentration-dependent dynamics of TF footprints, linked peaks to putative target genes, and elucidated rewiring of cis-regulatory networks. Our models reveal that early in reprogramming, OSK, at supraphysiological concentrations, rapidly open transient regulatory elements by occupying non-canonical low-affinity binding sites. As OSK concentration falls, the accessibility of these transient elements decays as a function of motif affinity. We find that these OSK-dependent transient elements sequester the somatic TF AP-1. This redistribution is strongly associated with the silencing of fibroblast-specific genes within individual nuclei. Together, our integrated single-cell resource and models reveal insights into the cis-regulatory code of reprogramming at unprecedented resolution. We establish a quantitative, predictive framework that links TF stoichiometry, motif syntax, and somatic silencing to provide new perspectives on the control of cell identity by TFs during fate transitions.

## Introduction

Cell state transitions proceed through a cascade of chromatin remodeling events and accompanying transcriptomic changes driven by cooperative and competitive binding of transcription factors (TFs) to complex syntax of sequence motifs encoded in cis-regulatory elements. Regulatory control of these cell state transitions can be systematically studied by perturbing the concentration of one or more critical lineage-defining TFs in a starting cell type and profiling molecular and phenotypic properties of perturbed cell populations over time^1^.

Through the course of these transitions, the stoichiometry of TFs can change quantitatively and their concentrations can span many orders of magnitude. How TF concentration dynamics of multiple TFs quantitatively orchestrate chromatin accessibility and concomitant target gene expression remains incompletely understood.

Reprogramming of differentiated cells into induced pluripotent stem (iPSCs) cells by ectopic expression of the Yamanaka TFs *POU5F1* (*OCT4*), *SOX2*, *KLF4* and *MYC* (OSKM) offers an ideal system to explore this interplay^2^. The cooperative action of this compact set of four TFs, which includes potent pioneers (*OCT4* and *SOX2*), provides a tractable opportunity to study TF stoichiometry and concentration dependent temporal dynamics of chromatin accessibility and the resulting activation and silencing of diverse gene expression programs.

Pioneering studies characterizing molecular features of fibroblast reprogramming have shown that overexpression of OSKM in somatic cells leads to extensive opening of closed chromatin, initiating multiple trajectories of cell state transitions, some of which culminate in successfully reprogrammed iPSCs^3–5^. Gene expression and chromatin profiling across human and mouse fibroblast reprogramming time courses show that somatic genes and regulatory elements are silenced early, while pluripotency-associated genes and regulatory elements are progressively activated in phases over 3 to 4 weeks^5–13^. Complementary approaches have demonstrated that the stoichiometry of ectopic OSKM at initiation and throughout the overexpression time course are critical determinants of the initiation, progression and efficiency of reprogramming^14–19^.

However, a quantitative framework connecting dynamic OSKM concentration, cis-regulatory syntax of their target sites, the resulting chromatin and gene expression state transitions, and ultimately reprogramming outcomes, has been lacking.

Recently, neural networks have emerged as state-of-the-art predictive models of regulatory DNA^20–25^. These models are trained to accurately map regulatory DNA sequences to associated experimental profiles of TF binding, chromatin accessibility, histone marks and gene expression in cellular contexts of interest. The models learn hierarchical layers of *de novo* sequence pattern detectors that can encode sequence motifs and their higher-order syntax.

Interpretation of these models has revealed novel insights into the cis-regulatory code of TF binding including sequence preferences and affinity landscapes of individual TFs^24,26^, motif-syntax mediated TF cooperativity^24,27^, and effects of sequence variation and repeats^20,22,26,28^. While neural networks have been used to dissect the sequence basis of chromatin accessibility of diverse cell types^29–32^, they have yet to be used to decipher regulatory dynamics during cellular reprogramming.

In this study, we present an interpretable deep learning framework applied to a densely-sampled, single-cell multi-omics dataset tracing OSKM induced reprogramming of human fibroblasts to iPSCs that reveals the interplay between combinatorial TF stoichiometry, motif affinity and syntax in determining cell fate. We discover that supraphysiological OSKM concentrations create transient regulatory elements absent from physiological cell states, which open early in the time course via occupation of non-canonical, low-affinity binding sites, and decay at rates determined by motif affinity as factor levels decline. These transient elements sequester the somatic factor AP-1 away from their endogenous targets, providing direct single-cell evidence for a competitive “repression-by-theft” mechanism that silences fibroblast gene programs and enables successful reprogramming. Our framework establishes a general paradigm for deciphering dynamic cis-regulatory codes that control cell fate from single-cell data.

### Single-cell transcriptomes and chromatin profiles of fibroblast reprogramming reveal diverse cell fate trajectories with continuous cell state transitions

To map regulatory and transcriptional cell states during human fibroblast reprogramming at high resolution, we overexpressed the reprogramming factors OSKM using a non-integrative Sendai virus system in primary dermal fibroblasts. We harvested cells at 9 time points and processed them for scRNA-seq and scATAC-seq (**Fig 1a**, **Methods**). The scATAC-seq data yielded 62,599 cells that clustered into 15 cell states (C1-15) (**Fig 1b**, **Fig X1a**, **Methods**). The initial day 0 population of fibroblasts and final population of iPSCs formed well separated clusters. On the other hand, samples from days 2-14 were composed of varying proportions of cells from different transient sub-populations, suggesting a continuum of cell states across the reprogramming time course (**Fig 1b,c**). We identified 525,835 scATAC-seq peaks representing putative cis-regulatory elements over all 15 cell states. The scRNA-seq data yielded 59,378 cells (**Fig 1d**, **Fig X1b**, **Methods**). The chromatin and expression landscapes across reprogramming were both consistent with an early loss of somatic identity and gradual gain of pluripotent identity^8^ (**Fig S1a,b,c, Supplementary Note**).

**Fig 1:**
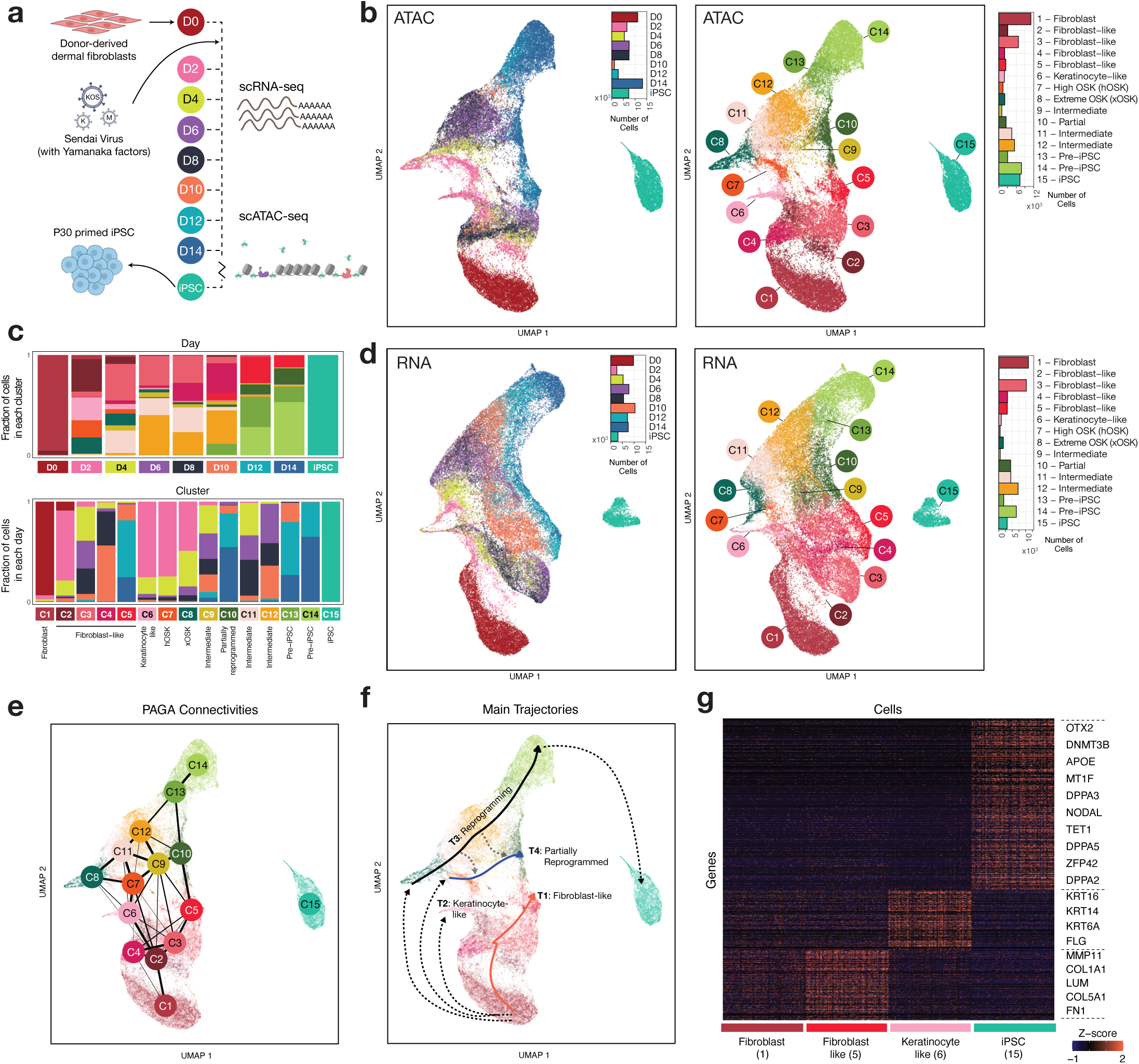
Single-cell chromatin and transcriptomic profiling of fibroblast reprogramming. **a)** Schematic of experimental design **b)** UMAP of single-cell ATAC-seq cells labeled by timepoint (left) and cell state identity (right) **c)** Barplots showing cluster-wise composition of scATAC-seq cells in each day (top), and day-wise composition of each cluster (bottom). The day-wise plots are normalized to account for differences in the total number of cells from each day. **d)** UMAP of single-cell RNA-seq cells labeled by timepoint (left) and cell state identity (right) **e)** PAGA connectivity graph derived from scATAC-seq data **f)** Key trajectories of cells overlaid on scATAC-seq UMAP **g)** Marker genes for the start state (Fibroblast) and multiple putative end states

We aligned cells from the scRNA-seq and scATAC-seq experiments in a joint CCA embedding. We transferred scATAC-seq derived cell state labels to cells in the RNA compendium and identified 156,016 putative regulatory peak-gene associations (**Fig S1d-g, Table S1, Methods, Supplementary Note**).

Next, we used the Partition-based graph abstraction (PAGA) method and diffusion based pseudotime inference on the scATAC-seq data to identify four main trajectories of cell state transitions (**Fig 1e,f,g, Fig S1h,i**, **Fig X2a-d, Methods, Supplementary Note**) ^33,34^. Two off-target trajectories terminate in stalled fibroblast-like cells (T1) and keratinocyte-like cells (T2), a primary reprogramming trajectory terminates in iPSC cells (T3), and a parallel trajectory ends in partially reprogrammed cells at day 14 (T4) .

We devised a new method to deconvolve endogenous and exogenous Sendai transcripts of OSKM (**Fig 2a**, **Methods**). Day 2 cells in clusters C7 and C8 that appear to initiate reprogramming, show high and extreme levels of Sendai OSK expression relative to endogenous OSK expression in iPSCs (**Fig 2a**) and were hence labeled as High OSK (hOSK) and Extreme OSK (xOSK) respectively. These states were disconnected from the initial fibroblast population (C1), suggesting that ectopic overexpression of reprogramming factors dramatically shifts the chromatin and transcriptomic landscapes and lends an appearance of cells “teleporting” early in the time course^4^. Cells in the reprogramming trajectory (T3) starting from the xOSK state (C8) proceed through intermediate states (C11, 12) and Pre-iPSC states (C13, 14) before terminating in the iPSC state (C15) after 30 rounds of passaging. The iPSC cells (C15) formed a disconnected cluster, suggesting a substantially altered state relative to Pre-iPSCs (C13, 14).

**Fig 2:**
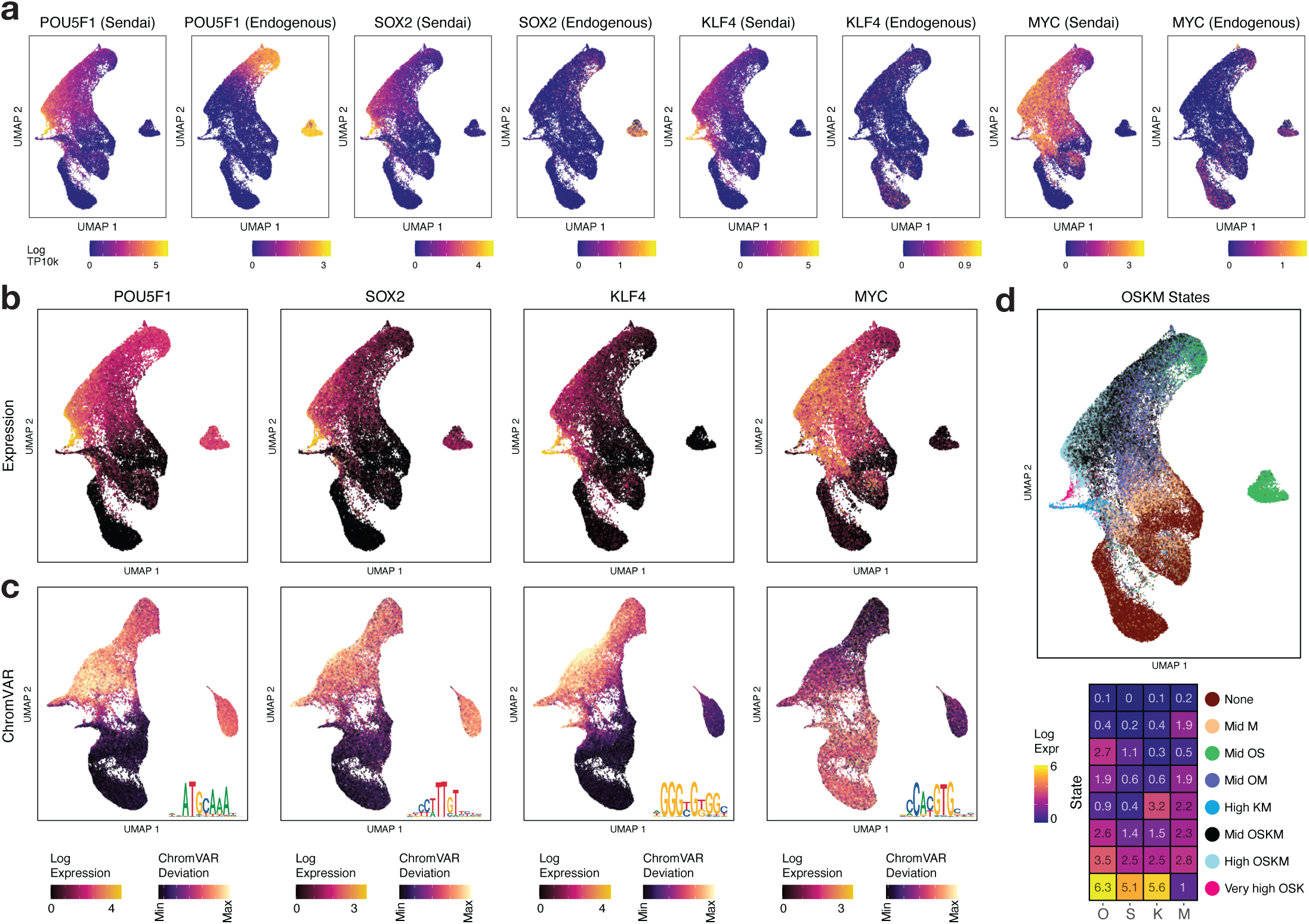
Stoichiometry and temporal dynamics of OSKM expression. **a)** Estimated exogenous (Sendai-derived) and endogenous expression of OSKM **b)** Total expression of OSKM factors overlaid on scRNA-seq UMAP **c)** ChromVAR deviation scores of OSKM motifs overlaid on scATAC-seq UMAP. Motif shown as inset. **d)** States derived from k-means clustering of OSKM expression: scRNA-seq cells labeled by OSKM state (top) and centroid expression values for each state (bottom)

In contrast, cells along the partially reprogrammed trajectory (T4) starting from the hOSK state (C7) proceed through intermediate state C9 and finally end in the partially reprogrammed state (C10), which fail to activate endogenous *OCT4* expression, in stark contrast to Pre-iPSC cells (**Fig 2a**). Furthermore, transgene expression levels were higher in states along the reprogramming trajectory (T3) compared to the partially reprogrammed trajectory (T4), suggesting that some cells fall potentially off the primary reprogramming trajectory (T3) into the partially reprogrammed trajectory (T4) (**Fig 1f** gray dotted arrows, **Fig S1j**, **Methods**). This is consistent with the observation that premature withdrawal of OSKM expression can stall the progress of cells in a partially reprogrammed state^35^. These results highlight the critical role of TF dosage in determining cell fate and motivates a deeper investigation into the underlying stoichiometric drivers.

### Stoichiometry and temporal dynamics of OSKM expression drive diverse cell fate trajectories

The absolute and relative stoichiometry of ectopic OSK at initiation and over the overexpression time course have profound effects on the progression and efficiency of reprogramming^14–18^. In our system, we reprogram cells using 3 vectors: *KLF4*, *MYC*, and a polycistronic KOS vector (**Fig 1a**). The inherent stochasticity of transgene expression and its temporal dynamics sets up a natural experiment that allows us to dissect cell states and fates associated with different combinations and doses of the reprogramming TFs over the course of reprogramming.

We observed systematic differences in temporal dynamics of total scRNA-seq expression levels of OSKM as well as their deconvolved endogenous and exogenous levels across cell state transitions (**Fig 2a,b, Fig S2a**, **Methods**). The broad patterns of variation of total OSKM expression were also reflected in their ChromVAR deviation motif scores (**Fig 2c, Fig S2b,c,d Methods**)^36^. The expression and ChromVAR deviation scores across cells correlated strongly for OSK but not *MYC* (**Fig S2c**), which is likely due to *MYC*’s inability to directly initiate chromatin accessibility in contrast to the *OSK* pioneer TFs^3,7^.

To quantitatively understand how the combinatorial expression of OSKM varied across the reprogramming time course, we clustered cells into 8 states based on single-cell expression levels of OSKM (**Fig 2d, Fig S2a,b**, **Methods**). Fibroblast-like cells in clusters C2-C5 along trajectory T1 expressed either low levels of OSKM or elevated levels of *MYC* only. Keratinocyte-like cells (C6) expressed high levels of KM but not OS.

Cells that seemed to initiate reprogramming at day 2 express higher levels of OSKM, with *OCT4* expression levels exceeding those of the other factors (**Fig S2a,b**). Day 2 cells in the xOSK state expressed *OCT4* 4-fold higher and *SOX2* 8-fold higher relative to iPSC levels (xOSK: *OCT4*: 7217 TPM, *SOX2*: 2194 TPM; iPSC: *OCT4*: 1740 TPM, *SOX2*: 271 TPM) (**Fig S2a**). OSKM transgene expression levels gradually attenuated over the time course, owing to dilution of Sendai virus vectors^37^. Compared to day 2 cells in xOSK state, day 10 cells in the Intermediate state C12 along the reprogramming trajectory T3 were down nearly 10-fold for OSK (**Fig S2a**) and the final population of iPSCs was transgene-free. Transgene expression levels were higher along states on the primary reprogramming trajectory (T3) compared to the partially reprogrammed trajectory (T4) (**Supplementary Note**).

In summary, we observe that cells with reprogramming potential transiently express supraphysiological levels of OSKM, supporting the observation that sustained high expression of OSKM may be critical for successful reprogramming^19^. Together, these results establish a direct link between OSKM stoichiometry and reprogramming trajectories.

### Base-resolution deep learning models reveal cis-regulatory sequence codes of cell-state resolved chromatin accessibility landscapes

A quantitative understanding of how cis-regulatory sequence influences TF binding and chromatin accessibility dynamics across diverse reprogramming fates has been elusive due to lack of high-resolution data and the combinatorial complexity of the cis-regulatory code. To decipher the cis-regulatory sequence code of accessible regulatory elements across the reprogramming time course, we trained convolutional neural network models called ChromBPNet^38^. ChromBPNet learns sequence pattern detectors akin to TF binding motifs and their higher-order organizational syntax from DNA sequence to predict base-resolution, pseudo-bulk scATAC-seq coverage profiles using ∼2kb local sequence context around scATAC-seq peaks and in background regions from each cell state (**Fig 3a**, **Methods**).

**Fig 3:**
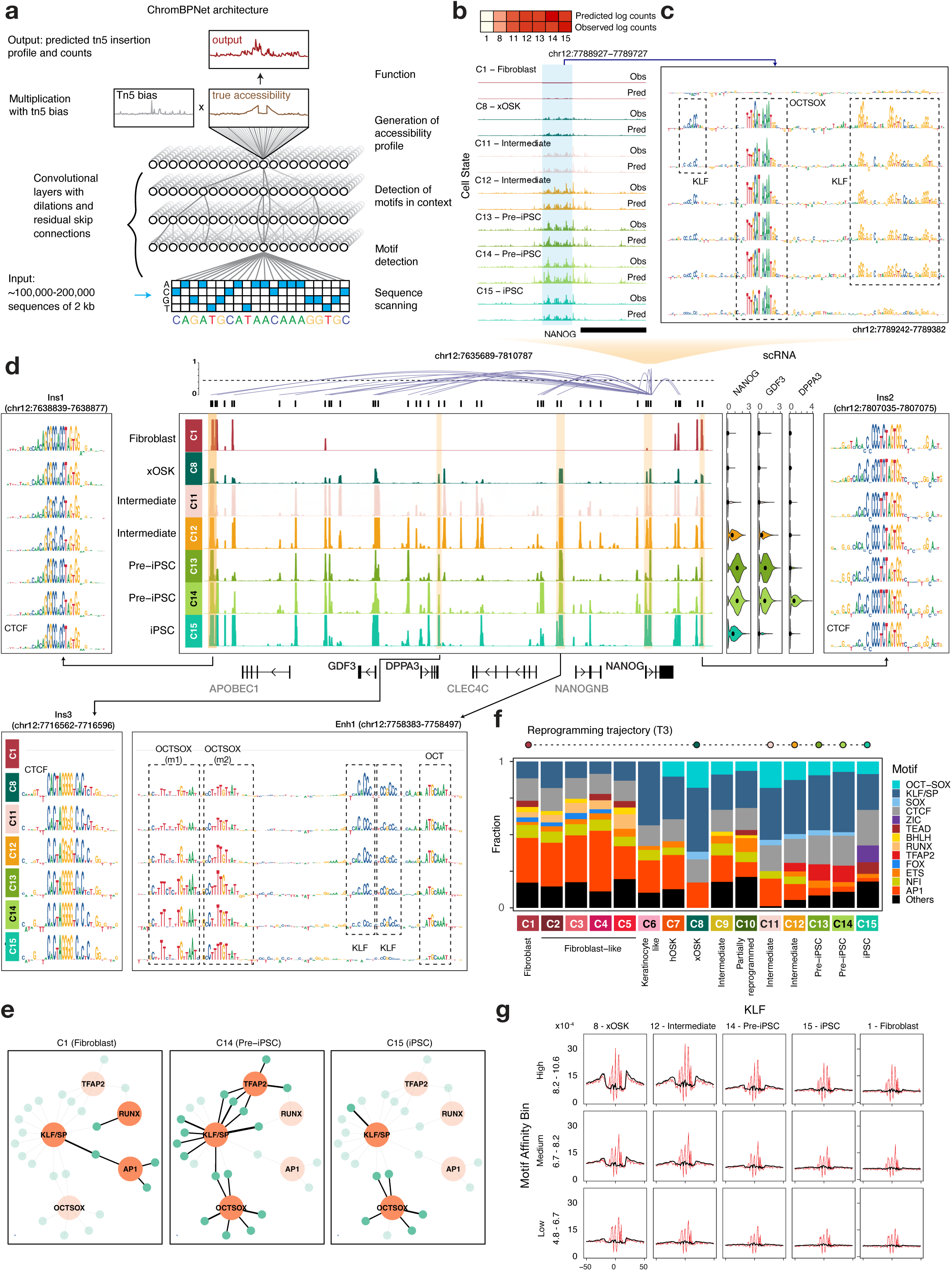
Base-resolution deep learning models to learn cell-state specific cis-regulatory sequence code. **a)** ChromBPNet model architecture: A single ChromBPNet model predicts the base-resolution pseudo-bulk scATAC-seq signal for a given cell state for a 2kb input sequence, while correcting for Tn5 sequence bias **b)** Normalized counts prediction of ChromBPNet models for each of the cell states along the primary reprogramming trajectory (T3) at the *NANOG* promoter (chr12:7788327-7790327) (top) and per base predictions over the central 1kb (bottom) **c)** Model-derived cell state-specific contribution scores directly upstream of the *NANOG* promoter. **d)** Pseudo-bulk scATAC-seq read pileup in the 176kb domain containing *NANOG* (chr12:7635000-7811000), along with expression of *NANOG*, *GDF3* and *DPPA3*, and peak-gene links for *NANOG* (center). Contribution reads for selected peaks within the domain. Contribution scores are shown only for cell states for which the locus was called as a peak. **e)** TF-to-gene network of NANOG: motifs are represented by orange nodes and peaks by blue nodes. Motif nodes are on when the motif is learned by the model for that cell type, and peak nodes are active if the peak is accessible in the cell type. An edge is active if an instance of the motif is found in the peak. **f)** For each cell state, the relative proportion of predictive instances of prominent motifs out of all predictive instances. **g)** Virtual footprinting for KLF obtained by inserting instances of KLF stratified by log-odds scores into random background sequences and averaging predicted profile probability distributions with (red) and without (black) bias for each cell state. States are ordered by decreasing KLF4 concentration.

ChromBPNet jointly models the total Tn5 insertion counts and their distribution (shape of the profiles) across these windows at single base resolution, as a function of the underlying sequence, while regressing out biases due to Tn5’s intrinsic sequence preference. We obtained high and stable Pearson correlation between total predicted and observed Tn5 insertion coverage in test regions across 10 cross-validation folds and cell states (**Fig S3a,b**). The observed and predicted base-resolution distributions of Tn5 insertions (shapes of the profiles) in test peak regions were also concordant (**Fig S3c,d**). For example, ChromBPNet models accurately recover the changes in magnitude and shape of accessibility profiles at the *NANOG* promoter (chr12:7788327-7790327) as it gains accessibility across the reprogramming time course (**Fig 3b**).

We used the DeepLIFT/DeepSHAP algorithm to derive the quantitative contribution of each base in any regulatory sequence of interest to its corresponding predicted accessibility profile from each cell-state-specific model (**Fig 3c**)^39,40^. At the *NANOG* promoter sequence, DeepLIFT reveals bases overlapping an evolutionarily conserved, high-affinity OCT-SOX binding motif gaining predictive contribution in the earliest time point after induction (xOSK state) and retaining high contribution through the time course, including the iPSC state^41^. In contrast, a constellation of adjacent KLF binding motifs first engaged in the xOSK state, show diminishing contributions through the time course with a complete loss in the iPSC state, which corresponds with the lack of *KLF4* expression in fibroblasts and iPSCs (**Fig S2a,b**). Hence, contribution scores derived from cell-state-specific ChromBPNet models can be used to infer putative bound TF motif instances that influence chromatin accessibility and also track putative TF binding occupancy dynamics across the time course in terms of motif contributions to chromatin accessibility.

We used our model-derived contribution scores to further explore the sequence features regulating chromatin accessibility dynamics of all scATAC-seq peaks in the ∼176kb domain containing the *NANOG* gene (**Fig 3d, Fig S4a**). This domain is largely inaccessible in fibroblasts and is flanked by two constitutively accessible peaks (Ins1 and Ins2) containing CTCF motifs with ubiquitously high contribution scores, which overlap loop anchors in micro-C contact maps in fibroblasts and ESCs (**Fig S4b**) ^42^. A putative enhancer element (Enh1) ∼30 kb upstream of the *NANOG* TSS is one of the earliest to gain accessibility in the xOSK state as a likely early target of OSK over expression. The contribution scores support this hypothesis by highlighting a constellation of OCT-SOX, KLF and OCT motifs which are predicted to open chromatin post-induction (xOSK state) and maintain accessibility throughout the time course post-induction. The KLF motifs once again show diminishing contributions over the time course with no contributions in the iPSC state (**Fig S2a**).

We constructed cell-state-specific TF-to-gene (TF2G) networks regulating *NANOG* by summarizing the TF binding sites with high contribution scores in all dynamic peaks linked to *NANOG* (**Fig 3e**, **S4c Methods**). These networks implicate OSK and somatic TFs RUNX and AP-1 driving chromatin dynamics early in reprogramming. *NANOG* expression is highest in the Pre-iPSC states (C13, 14) which correspond to extensive co-occupation of putative enhancers by OSK and TFAP2. TF2G networks of *FN1* and *JUN* genes similarly demonstrate how ChromBPNet enables intricate, multi-scale dissection of the cell-type resolved, dynamic cis-regulatory sequence code at the resolution of individual base-pairs, elements and regulatory domains of genes (**Fig S5a,b,c, Supplementary Note**).

Next, we used the ChromBPNet models to derive base-resolution contribution scores for all accessible peaks in each cell state. While high contribution sequence patterns often resembled canonical motifs of TFs, we occasionally observed significant deviations. For example, a high contribution score motif instance (m2) in the earliest *NANOG* enhancer (Enh1) resembles a partial version of known OCT-SOX motifs and scores poorly as a sequence match to any known motif^43^ (**Fig 3d**). This highlights the unique ability of our predictive models to identify functionally important non-canonical sites that would otherwise be missed. We used the TF-MoDISco algorithm for *de novo* discovery of consolidated motif patterns from the sequence contribution score profiles of all peaks^44^. TF-MoDISco identified 30 non-redundant, *de novo* motifs across all 15 cell states (**Fig S6a**). We scanned all peaks to obtain a comprehensive annotation of predictive motif instances across the genome and summarize variation in the number of predicted motif instances across the cell states (**Fig 3f**, **Methods**). In Fibroblasts (C1), AP-1 and CTCF are the most prominent predictive motifs. Post OSKM induction, the accessible chromatin landscape is dominated by OCT-SOX, SOX and KLF motifs in the xOSK (C8) state which account for >50% of all predictive motif instances recovered from peaks. In successive states, the fraction of OSK motifs reduces in tandem with a withdrawal of OSK expression levels. TFAP2 motifs transiently drive accessibility in Pre-iPSC cells (C13, C14). ZIC and TEAD motifs gain prominence in the iPSC state (C15), while CTCF motifs increase to account for nearly 24% of all motif instances. These results suggest that chromatin dynamics in reprogramming is encoded in a compact, combinatorial and dynamic TF motif lexicon.

### Transcription factor concentration and motif affinity influence scATAC-seq footprints across reprogramming

Analyzing Tn5 transposition pile-ups at nucleotide resolution can reveal TF “footprints” that provide support for TF occupancy at motif instances^45^. The characteristic depth and shape of TF footprint profiles at canonical motif instances have been previously shown to be a function of residence time of the TF^45^. However, the interplay between TF concentration and footprint morphology has not been explored previously.

We can use ChromBPNet models to correct for Tn5 bias and predict bias-corrected ATAC-seq profiles for any genomic or synthetic sequence. For example, at the CTCF instance Ins2 in the *NANOG* locus, the uncorrected predicted scATAC-seq profiles closely match observed scATAC-seq profiles across all cell states and show a strong footprint over the CTCF motif (**Fig S7a, Fig 3d**). However, the bias-corrected predicted profiles differ substantially from the observed and uncorrected profiles, highlighting a deeper, narrower footprint with increased protection from Tn5 transposition right over the CTCF motif.

We next performed “in-silico marginal footprinting” to uncover the influence of TF concentration on predicted TF footprints. We embedded each motif in a library of inaccessible GC-matched background sequences randomly sampled from the genome, predicted uncorrected and bias-corrected scATAC-seq profiles for the library using ChromBPNet and averaged the profiles over the entire library to derive marginal uncorrected and bias-corrected TF footprint profiles (**Methods**). We predicted a comprehensive catalog of marginal footprints for all 30 TF-MoDISco motifs over a range of motif affinities binned into 3 strata (high, medium and low affinity), using ChromBPNet models from the 15 cell states which exhibit dramatic changes in TF concentrations across the reprogramming time course (**Fig X3, Fig S7b**).

For each affinity stratum of embedded KLF motifs, the strength of marginal footprints tracks expression of *KLF4* across cell states (**Fig 3g**). Similarly, for each cell state, marginal KLF footprints get stronger with increasing motif affinity. The footprints are strongest in the xOSK cell state. Even the weakest affinity motif stratum shows detectable footprints. In contrast, the weakest affinity motifs do not generate footprints in Pre-iPSCs at intermediate levels (∼80 TPM) of *KLF4* expression. iPSCs and fibroblasts, which have trace levels of *KLF4* expression, do not show footprints across the affinity spectrum including the strongest motif instances.

ChromBPNet models predict shallow but detectable marginal footprints at SOX and OCT-SOX motifs exclusively in the hOSK and xOSK states (**Fig S7c,d**). These TFs typically lack detectable DNase-I footprints at occupied motif instances in ChIP-seq peaks in mESCs at physiological TF concentrations, presumably due to their short residence times^45,46^. However, ChromBPNet models suggest that at very high concentrations, even these low-residence time TFs can in fact generate footprints. The somatic TF AP-1 also shows a strong relationship between marginal footprint strength, concentration and motif affinity (**Fig S7e**). The predicted in-silico marginal footprints are consistent with average footprints derived from observed Tn5 insertions over motif instances in accessible peaks (**Fig S7f,g,h**).

These results reinforce the biophysical principle that both TF concentration and motif affinity influence TF occupancy which is often directly reflected in morphology of TF footprints in ATAC-seq profiles after bias correction. Our models provide, for the first time, an avenue to predict and dissect this relationship at unprecedented resolution and fidelity across dynamic cell states.

### Peak sets of coordinated chromatin dynamics are regulated by distinct dynamic regulatory syntax

Overexpression of OSKM dramatically reconfigures the chromatin landscape of fibroblasts (**Fig S8a**). To identify modules of open chromatin regions with synchronized dynamics, we clustered the 525,835 scATAC-seq peaks into 20 peak sets across 7 major archetypes of distinct patterns of accessibility (**Fig 4a, Fig S8b, Methods**). For each peak set, we identified putative target genes, performed gene set enrichment analysis, and quantified global TF motif activity (**Fig 4b,c,d, Table S2, Methods**).

**Fig 4:**
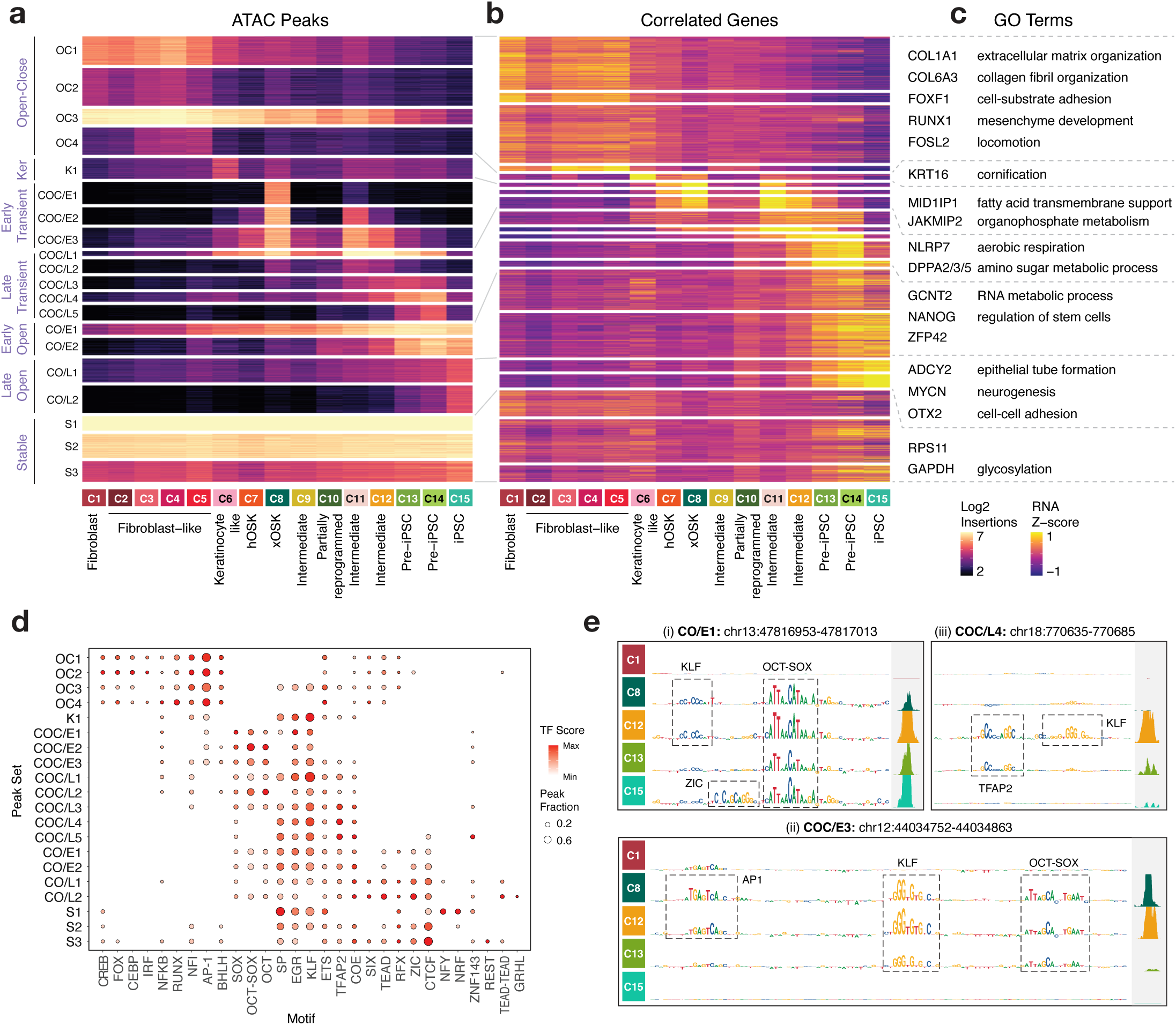
Monotonic and transient chromatin peak sets. **a)** Normalized accessibility of peaks across cell states grouped into 20 peak sets, further grouped into 7 major archetypes **b)** Normalized gene expression of genes linked to each of the peaks in each peak set **c)** Representative genes and gene set enrichment terms summarized for each peak set archetype **d)** Motif activity scores and abundances for each peak set **e)** ChromBPNet contribution scores and observed accessibility signal at 3 representative loci

As expected, the Open-Close peak sets (OC1-4) are enriched for somatic TFs such as AP-1, ATF, and CEBP and are linked to the regulation of fibroblast-specific genes. Similarly, Close-Open peak sets consist of iPSC-specific peaks that gained accessibility either early (CO/E1-2) or late (CO/L1-2) during reprogramming. CO peaks are involved in stem cell regulation, RNA metabolic process, cell-cell adhesion, and neurogenesis processes, consistent with previous analyses^6^. They are primarily regulated by OCT-SOX and KLF motifs as well as CTCF, TEAD, and ZIC motifs (**Fig 4d, e (i)**).

Beyond peaks with monotonic accessibility patterns, nearly 40% of all dynamic peaks along the primary reprogramming trajectory (T3) are transient (**Supplementary Note**). Early Transient peaks (COC/E1-3) are closed in fibroblasts but open at the first time point after OSKM induction in the xOSK (C8) state, before closing eventually in iPSCs. COC/E peaks are enriched primarily for OSK motifs, suggesting that they arise as an immediate consequence of OSKM overexpression. In addition, predictive instances of AP-1 motifs also feature in Early Transient peaks, consistent with previously observed redistribution of somatic TFs to transient sites measured by ChIP-seq in mouse (**Fig 4d, e(ii)**)^5,7,9^. Since *OCT4* and *SOX2* are constitutively expressed in all cell states from xOSK through iPSCs, and *KLF4* is also expressed till day 14 Pre-iPSC (C14) cells (**Fig S2a**), the loss of COC/E peaks a few days after their opening strongly hints at stoichiometry-dependent action. Pseudotime analysis of TF expression and motif activity along the reprogramming trajectory (T3) further suggests expression dependent binding activity of OSK and AP-1 (**Fig S8c,d**, **Methods**). Notably, 1913 genes linked with all Early Transient peak sets (COC/E) lack coherent functional annotation based on gene-ontology enrichment, consistent with previous studies (**Table S2, Fig S9a**)^6^. In subsequent sections, we perform a detailed exploration of the regulatory syntax and potential role of this large set of enigmatic transiently active regulatory elements in reprogramming.

### Early Transient peaks are unique to reprogramming and not found in differentiated cell types and tissues

A key question is whether these Early Transient peaks (COC/E) represent a normal, transient developmental state, or are a unique artifact of OSKM reprogramming. To distinguish between these possibilities, we first checked whether they overlapped accessible sites in a reference DNase I hypersensitive sites (DHS) atlas across 438 well characterized cell-lines, primary cells and tissues^47^ (**Fig 5a**, **Methods**). Nearly half of the peaks in Early Transient peak sets COC/E1 and COC/E2 are absent in all biosamples from the DHS index. In contrast, <5% of the peaks in Open-Close (OC1-3) and Stable peaks (S1-3) sets are absent in the DHS Index. The stark depletion of the early transient peaks in diverse cell types and tissues suggests these peaks may be a unique feature of our reprogramming time course.

**Fig 5:**
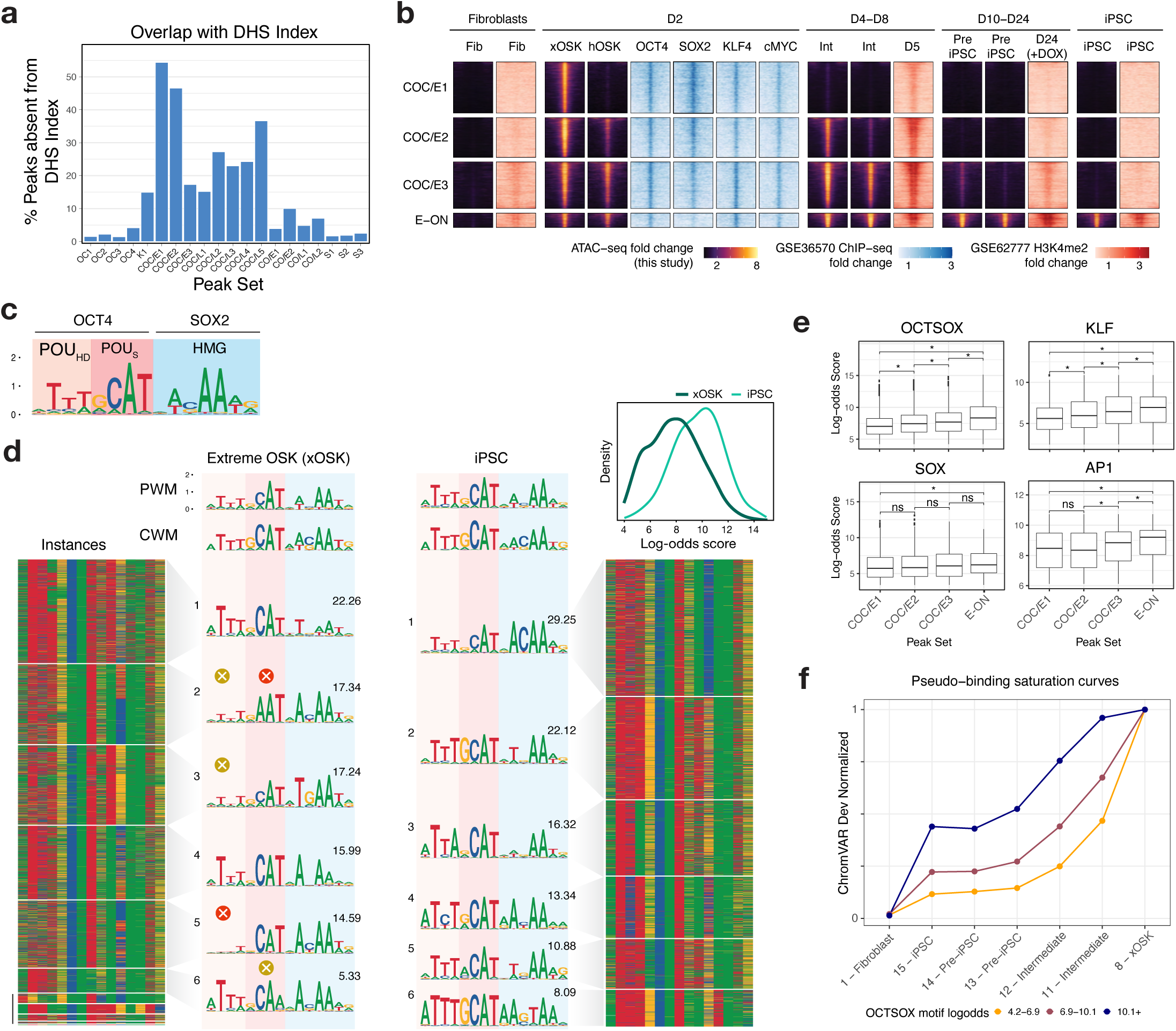
Characterization of Transient peaks sets. **a)** Fraction of peaks in each peak set that are not present in a reference DHS Index of 438 biosamples **b)** Integration of scATAC-seq data with day 2 OSKM ChIP-seq data (GSE36570) and H3K4me2 ChIP-seq from fibroblasts, days 5, 24 and iPSC (GSE62777). 2kb regions are shown for scATAC-seq and OSKM ChIP-seq, and 4kb regions for H3K4me2 ChIP-seq **c)** Canonical OCT-SOX motif from (Avsec et al. 2021), labeled with its binding domains **d)** TF-MoDISco OCT-SOX motif recovered in the xOSK and iPSC cell states. Each row in the heatmap is one genomic instance of the OCT-SOX motif retrieved by TF-MoDISco. The instances are clustered into sub-motifs separately for each cell state by TF-MoDISco, with the relative fraction of each sub-motif indicated above its PWM. The inset shows the distribution of the log-odds scores over motif instances in the two cell types. **e)** Box-plot of the distribution of log-odds scores for motif instances of OCT-SOX, KLF, SOX and AP1 motifs within COC/E1-3 and E-ON peak sets, * *p*-value < 0.001 **f)** Pseudo-binding saturation curve for OCT-SOX. Each line is one strata of peaks split based on the affinity of the OCT-SOX motif/s present in them. States are ordered in roughly increasing order of combined OCT4 and SOX2 gene expression.

To determine whether these early transient peaks are a general feature of reprogramming, we compared the Early Transient peak sets (COC/E1-3) and a contrastive set of peaks (E-ON) that opened early at Day 2 and remained accessible throughout our time course against OSKM ChIP-seq^3^ and H3K4me2 histone ChIP-seq^6^ (**Fig 5b**) data from two alternative reprogramming systems. Unlike our Sendai virus based reprogramming of dermal fibroblasts, Soufi et al. used doxycyline (dox)-inducible lentiviral transduction while Cacchiarelli et al. used secondary fibroblasts with constitutive human telomerase (hTERT) expression. All Early Transient and E-ON peak sets show strong enrichment of OSKM binding from ChIP-seq data at day 2 from Soufi et al. Moreover, H3K4me2 mirrors chromatin accessibility patterns of both peak sets across the time course^6^. Hence, the early transient peaks are a reproducible signature of multiple OSKM induced fibroblast reprogramming systems.

### OCT/SOX target low-affinity non-canonical motifs at high initial concentrations to open chromatin in Early Transient peaks

Next, we investigated cis and trans regulatory factors that could potentially explain the gain and eventual loss of Early Transient peaks. The mRNA levels of *OCT4* and *SOX2* are 4-fold and 8-fold higher in xOSK compared to iPSCs (C15), and OSK motifs disproportionately dominate the chromatin landscape in the xOSK state (**Fig 3f**). We suspected that supraphysiological concentration and stoichiometry of OS in the xOSK states could enable these pioneer factors to occupy low affinity, non-canonical binding sites resulting in esoteric landscapes of chromatin accessibility. Based on this hypothesis, we expected early transient peaks in the xOSK state to be enriched for non-canonical OCT-SOX binding sites.

To test this hypothesis, we first compared the *de novo* OCT-SOX motifs inferred by TF-MoDISco from ChromBPNet models of xOSK and iPSC states to a canonical OCT-SOX motif^24^. The TF-MoDISco OCT-SOX motifs from xOSK and iPSC states were similar to the canonical OCT-SOX heterodimer motif which consists of three parts - a consensus subsequence ATTT followed by GCAT which are recognized by OCT4’s POU-homeodomain (POU_HD_) and POU-specific (POU_S_) domains respectively, and a trailing ACAA subsequence that binds to the SOX2 HMG domain (**Fig 5c,d**)^48^. However, the xOSK-derived OCT-SOX motif showed subtle differences from the canonical motif in the two POU recognition subsequences. To obtain a deeper understanding of these differences, we identified all predictive motif instances of the xOSK and iPSC-derived TF-MoDISco OCT-SOX motifs from the xOSK and iPSC peaks respectively. We then computed the log-odds scores of the motif instances from xOSK and iPSCs against the canonical OCT-SOX motif as a surrogate measure of affinity. OCT-SOX motif instances from xOSK cells showed significantly lower log-odds scores compared to instances from iPSCs (**Fig 5d**, inset, Mann-Whitney *p*-value <2.2×10^-16^) (**Methods**), suggesting that the early transient peaks likely harbor low affinity OCT-SOX binding sites.

To reveal fine-resolution heterogeneity of OCT-SOX motifs in xOSK and iPSC peaks, we re-clustered the motif instances of the TF-MoDISco OCT-SOX motifs from each state into more granular sub-motifs (**Fig 5d**). All OCT-SOX sub-motifs from iPSCs are similar to the canonical OCT-SOX motif in terms of engaging both POU domains and SOX-HMG. In stark contrast, several OCT-SOX sub-motifs from xOSK differed substantially from the canonical OCT-SOX motif. xOSK sub-motifs 2 and 6 show distinct suboptimal matches to the canonical recognition sequence (GCAT) of the POU_S_ domain. Further, in OCT-SOX sub-motifs 2,3 and 5, the leading ATTT subsequence that binds the OCT4 POU_HD_ domain is greatly attenuated, suggesting that a partial POU_S_-HMG heterodimer motif in transient peaks may be sufficient to recruit the OCT4-SOX2 heterodimer at high concentrations in the xOSK state. A recent study supports our hypothesis by showing that OCT4-SOX2 heterodimers can in fact bind nucleosome-embedded instances of this exact partial motif and drive chromatin accessibility^49^. Our results also complement ChIP-seq experiments that show that OSK can individually target their partial monomeric motifs at nucleosome-embedded sites early during reprogramming^50^.

Thus, our models suggest that the early transient peaks in OSK-induced early reprogramming are a result of unique occupancy patterns of OCT4 and SOX2 pioneer factors at supraphysiological concentrations at non-canonical low-affinity heterodimeric binding sites.

### Duration of transience of chromatin accessibility correlates with motif affinity

The gradual loss of chromatin accessibility of early transient peaks from the xOSK state to iPSCs is associated with a smooth decrease in the concentration of the OSK factors. However, subsets of early transient peaks exhibit subtle differences in their decay rates across the time course (**Fig 4a**). The COC/E1 subset of peaks lose accessibility immediately after the xOSK state at day 2. COC/E2 peaks remain accessible longer until intermediate cell state 11 (day 8). COC/E3 peaks exhibit the slowest decay and are accessible through intermediate cell state 12 (day 8) until the Pre-iPSC state (day 10). The variable decay rates across these peak subsets could be due to subtle differences in the affinities of non-canonical OSK motif instances in each of these subsets. To test this hypothesis, we compared log-odds scores of instances of the OCT-SOX, KLF and SOX motifs across the three subsets of early transient peaks. We further contrasted these early transient peak subsets against a control set of peaks (E-ON) that also gain accessibility at day 2 but remain accessible throughout the trajectory and in iPSCs (**Fig 5e**, **Methods**). For all three motifs, we observed significant differences in log-odds distributions between the four peak sets, such that peak sets with faster decay rates contained motifs with lower surrogate motif affinity.

Next, we stratified peaks from all four peak sets into three levels of OCT-SOX motif affinity based on the highest scoring OCT-SOX motif instances in each peak. For peak sets corresponding to each affinity stratum, we compared their ChromVAR deviation motif scores to *OCT4*/*SOX2* expression levels across all the cell states in the primary reprogramming trajectory. These surrogate binding saturation curves for OCT-SOX not only show that accessibility decreases with TF concentration but also that the decay rates are faster for peaks with lower affinity OCT-SOX motifs (**Fig 5f**, **Methods**). Surrogate binding saturation curves for *KLF4* showed similar trends (**Fig S9c**).

Overall, our results suggest that early transient peaks gain accessibility in the xOSK state due to binding of OSK at supraphysiological concentrations to their motif instances across a wide range of binding affinity. The dynamic and variable rates of decay of accessibility across the time course can be attributed to progressive release of weaker affinity sites as TF concentrations gradually decline.

### Sequestration of AP-1 to transient sites is associated with somatic silencing at single-cell level

The functional role of early transient peaks is not well understood. Previous studies have suggested that transient peaks sequester somatic TFs such as AP-1 resulting in redistribution of somatic TFs away from fibroblast-specific peaks, thereby destabilizing the fibroblast chromatin landscape and promoting reprogramming^5,7^. Indeed, we find that the early transient peak sets are enriched for multiple predictive instances of somatic TF motifs, especially AP-1, alongside motifs of the OSK reprogramming factors (**Fig 4d,e (ii)**). Specifically, ∼10,500 predictive AP-1 motifs supported by discernable ATAC-seq footprints are identified in peaks that first gain accessibility in the xOSK state (**Fig 6a**), a majority (∼70%, i.e. ∼7,500) of which are found in the early transient peak sets. In contrast, ∼35,000 (51%) of the ∼69,000 predictive AP-1 motif instances found in fibroblast peaks are lost in the xOSK state.

**Fig 6:**
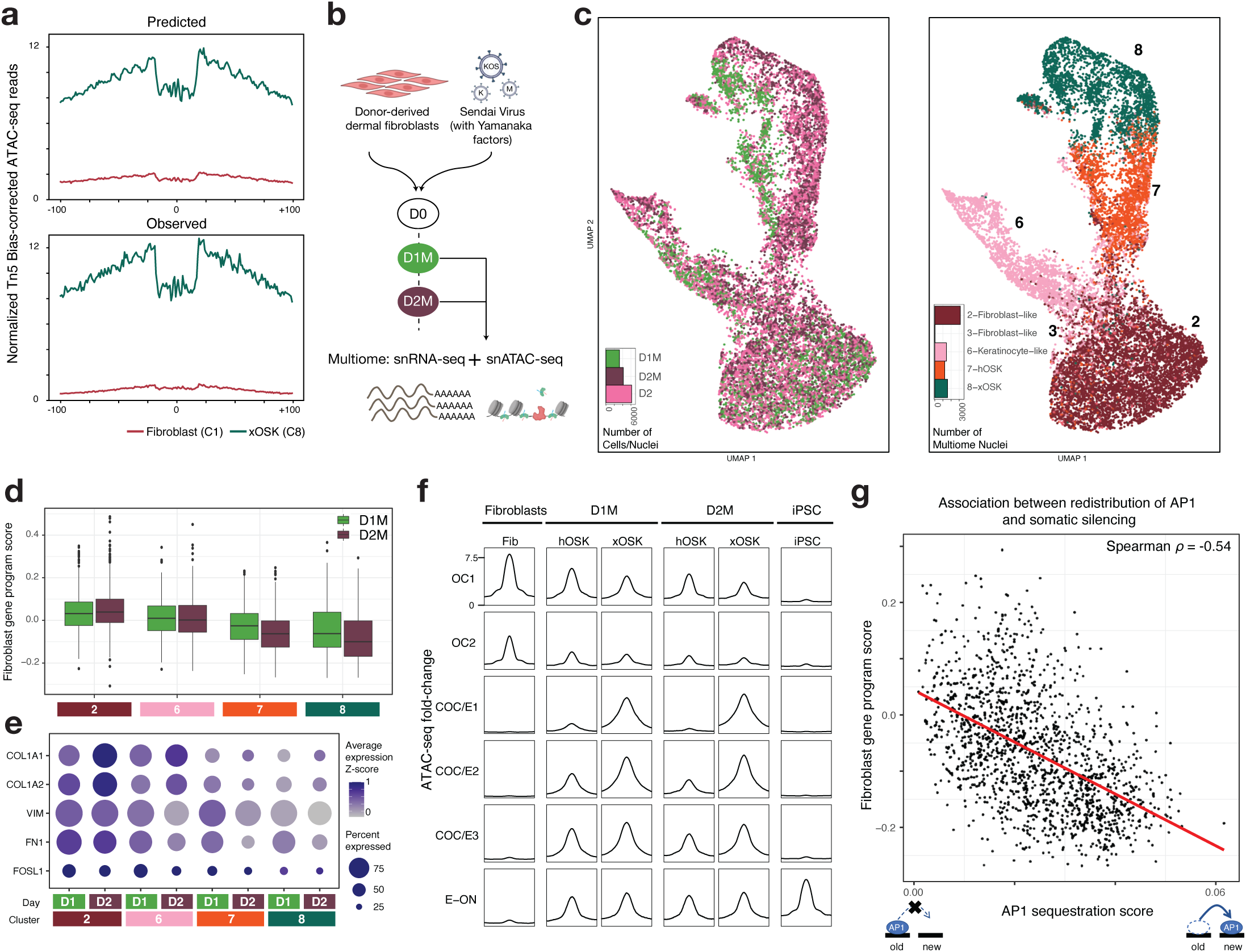
Single-nucleus multiome profiling highlights the role of Early Transient peaks in somatic repression. **a)** Predicted (top) and observed (bottom) Tn5 bias-corrected footprints over AP1 motifs within Early Transient peaks in the xOSK state **b)** Schematic of experimental design. M in D1M and D2M indicates Multiome. **c)** UMAP of multiome cells integrated with D2 scATAC-seq cells labeled by sample (left) and cell state (right) **d)** Box-plot of single-nucleus expression of fibroblast-specific genes stratified by day and cell state **e)** Single-nucleus expression of representative fibroblast-specific genes stratified by day and cell state **f)** scATAC-seq and snATAC-seq normalized read pileup at OC, COC/E and E-ON peak sets **g)** Scatter-plot of AP1 sequestration score versus snRNA-seq expression of fibroblast-specific genes. Each point is a single nucleus from Day 2 multiome data from hOSK and xOSK states.

We wondered whether there was an OSK dosage-dependent relationship between the displacement of somatic factors and repression of fibroblast gene sets. To study this phenomenon, we jointly profiled single-nucleus RNA-seq and ATAC-seq (sn-Multiome) at days 1 and 2 during the early stages of reprogramming (**Fig 6b**, **Methods**). We obtained 3217 nuclei from day 1 and 4161 nuclei from day 2 that passed quality control metrics (**Fig X4**, **Methods**). We annotated nuclei from the sn-Multiome data by transferring labels from our previously annotated reprogramming map after harmonization and integration using the ATAC-seq modality (**Fig 6c**, **S10a**). All early cell states (Fibroblast-like (C2, 3), Keratinocyte-like (C6), hOSK (C7), and xOSK (C8)) from the original time course were represented in the multiome data.

We first analyzed snRNA-seq expression of fibroblast-specific genes in each of the cell states in the multiome data to gauge the extent of somatic silencing in the first two days of reprogramming. We estimated a “fibroblast gene program score” for each nucleus as the average expression z-score of fibroblast-specific genes (**Fig 6d**, **Methods**). Compared to fibroblast-like cells (C2), all other cell states showed reduced expression of fibroblast-specific genes within the first two days, consistent with transcriptional silencing of somatic targets during early reprogramming^51^. The xOSK state exhibited the lowest fibroblast gene program score followed by hOSK, both of which exhibited reduced expression of fibroblast-specific genes at day 2 relative to day 1. Key fibroblast marker genes including *COL1A1* and *FN1* showed reduced expression as early as day 1 (**Fig 6e**).

Analogous analysis of the snATAC-seq data showed that chromatin accessibility at fibroblast-specific peaks in the OC1 and OC2 peak sets decreased rapidly after OSKM induction (**Fig 6f**). Simultaneously, early transient peaks in the COC/E1-3 peak sets gained accessibility, with higher accessibility in xOSK state mirroring our previous analysis of scATAC-seq data (**Fig 5b**). The rapid loss and gain of accessibility at distinct sets of regulatory elements agrees with previously observed enhancer dynamics as early as before a single cell division upon overexpression of Yamanaka factors in MEFs^51^.

We then used the multiome data to test for quantitative association between the degree of sequestration of AP-1 to early transient peaks and repression of fibroblast-specific genes. For each nucleus, we estimated an “AP-1 sequestration score” as the proportion of ATAC-seq reads falling in Early Transient peaks containing predictive AP-1 motif instances relative to total reads in all peaks (**Methods**). Focusing on nuclei from hOSK and xOSK cell states at day 2, we observed a striking negative correlation (Spearman ⍴= −0.54, *p*-value < 2.2×10^-16^) between the fibroblast gene program score and the AP-1 sequestration score over early transient peaks (**Fig 6g**). This negative association holds after including fibroblast-like nuclei and also for nuclei from day 1 (**Fig S10b,c**). In contrast, an analogously computed “AP-1 retention score” aggregated across fibroblast-specific peaks containing predictive AP-1 motifs was positively correlated with fibroblast gene program scores across nuclei from hOSK and xOSK cell states at day 2 (**Fig S10d**, Spearman ⍴=0.59, *p*-value < 2.2×10^-16^). These results provide the first direct, quantitative evidence linking sequestration of AP-1 to somatic gene silencing within the same cell.

Overexpression of OSKM thus has a quantitative effect on somatic silencing in the early stages of reprogramming, likely mediated by redistribution of somatic TFs to newly opened regulatory elements.

## Discussion

Our study provides a comprehensive understanding of the cis-regulatory code of cellular reprogramming by interpreting longitudinal single-cell multi-omics experiments through the lens of cell-state resolved deep learning models of regulatory DNA. By interrogating these models with powerful interpretation tools, we decipher quantitative relationships between dynamic OSKM concentration, motif affinity and syntax of their target sites, TF footprints, subsequent monotonic and transient chromatin state transitions, somatic program silencing, and ultimately successful pluripotency activation. We highlight six major contributions of our study that collectively establish a new paradigm for understanding factor-induced cell fate transitions.

### A high-resolution, single-cell, multi-omic resource of human fibroblast reprogramming

We present a unique resource of single-cell RNA-seq and ATAC-seq experiments representing high-resolution and temporally dense profiling of reprogramming intermediates, in contrast to past efforts that sorted reprogramming intermediates based on surface marker expression^4,6–10^. The single-nucleus multiome profiling enriches this atlas with an unprecedented view into the earliest stages of fibroblast reprogramming.

### State-of-the-art deep learning models decode the dynamic cis-regulatory code of reprogramming

This study represents the first application of deep learning models to scATAC-seq profiles in reprogramming. Our ChromBPNet models can predict cell-state resolved base-resolution ATAC-seq profiles with high accuracy from local sequence context, while providing state-of-the-art correction of confounding enzymatic biases, thereby revealing latent, de-noised TF footprints and the underlying predictive sequence syntax.

Our study specifically highlights the unique capability of ChromBPNet to unravel stoichiometry-dependent binding of TFs to non-canonical low-affinity binding sites. Classical motif annotation methods based on scanning sequences with oversimplified position weight matrix (PWM) motif models and arbitrary scoring thresholds struggle to reliably detect low affinity TF binding sites due to poor precision-recall tradeoffs. In contrast, ChromBPNet can learn complex *de novo* representations of sequence features and their higher-order syntax that are explicitly predictive of ATAC-seq profiles in each cell state. The model-derived contribution scores thus highlight base-pairs that quantitatively influence accessibility profiles without relying on PWM motif models of sequence affinity and arbitrary scoring thresholds. Our analyses show that the relative contribution of each TF to accessibility is influenced by motif affinity, TF concentration and the arrangement of other motifs (syntax) in the local context, which is reflected both in the contribution scores and the morphological properties of bias-corrected ATAC-seq footprints (**Fig 3g**).

We used the models to derive predictive TF motif syntax and associated TF footprints encoded in thousands of dynamic regulatory elements, and to elucidate the rewiring of cis-regulatory networks across diverse trajectories. Our framework enables estimation of the dynamic occupancy of TFs across cell states at motif instances traversing a wide affinity spectrum conditioned on TF concentration and their impact on quantitative changes in chromatin accessibility in the local sequence context of individual enhancers.

### Reprogramming traverses non-physiological regulatory landscapes

Chromatin accessibility and gene expression dynamics across the reprogramming time course reveal that OSKM combinatorially and quantitatively reconfigure the somatic fibroblast cell state. Upon induction, stochastic variation in exogenous OSKM stoichiometry results in initial diversification into a putative reprogramming trajectory, and off-target keratinocyte-like and fibroblast-like fates. In cells with supraphysiological overexpression of all factors, OSK rapidly dominate the chromatin landscape by extensively binding to and opening closed chromatin, while fibroblast-specific peaks close early (**Fig 7i-iii**). Newly accessible sites primarily contain OSK motifs, which unambiguously supports the view that OSK act as pioneer factors upon overexpression^8,9,49^.

**Fig 7:**
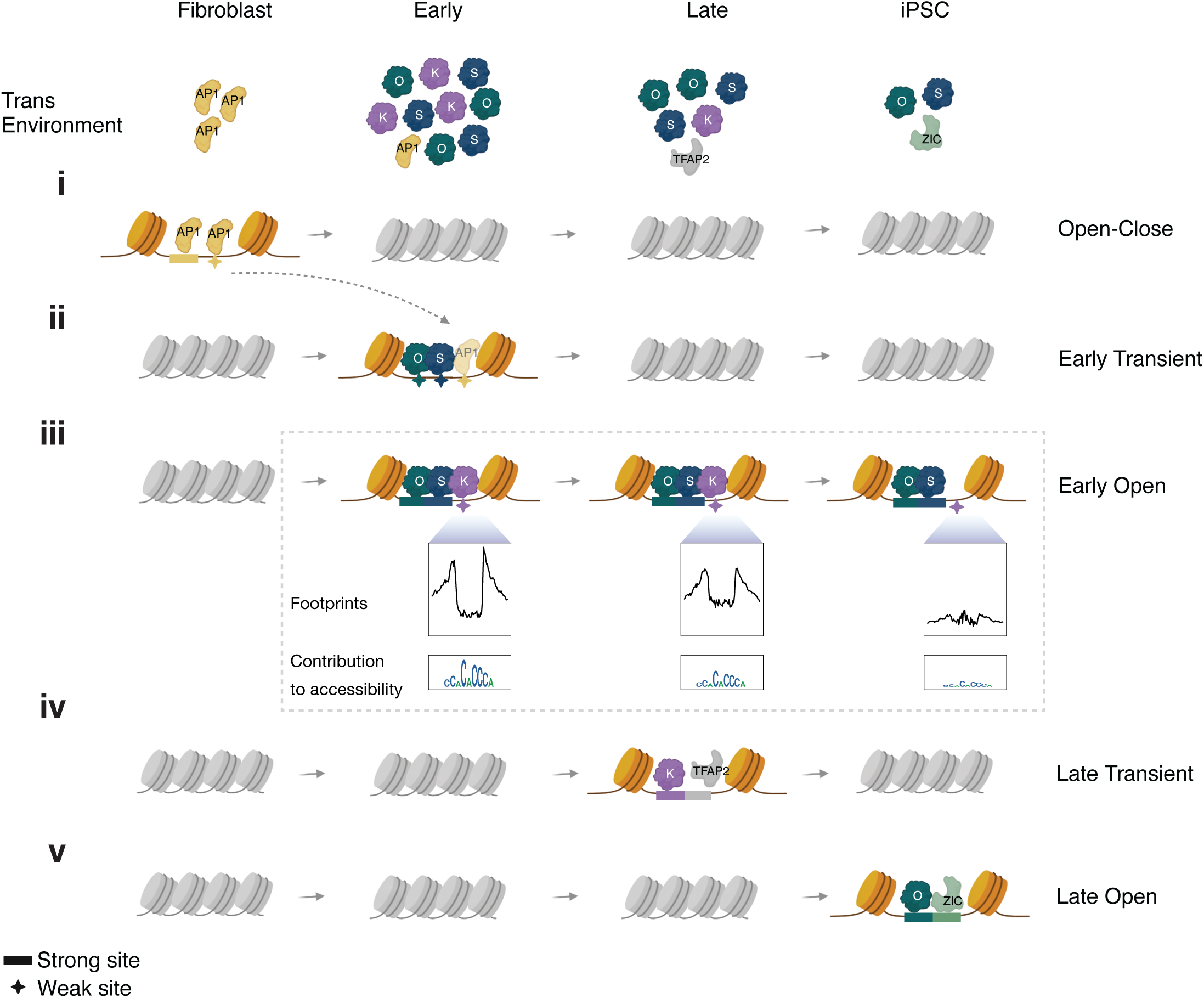
**A refined model of reprogramming adapted from (Chronis et al. 2017)**

Intriguingly, a majority of sites that open early are lost over the course of reprogramming, concomitant with a reduction in the expression levels of OSKM. We show that these Early Transient peaks are a reproducible signature of OSKM reprogramming and most of these peaks are exclusive to fibroblast reprogramming. The gene expression program driven by early transient peaks also does not map to any well characterized gene sets. These findings emphasize that reprogramming is not a simple reversal of development, and extreme OSKM expression in the initial phase pushes cells through a novel, non-physiological cell state.

### Supraphysiological TF concentrations enable occupation of low-affinity sites in transient peaks

Within Early Transient regulatory elements, we find pervasive evidence of non-canonical, low-affinity binding sites of OSK, including an excess of partial POU_S_-HMG binding motifs. These observations suggest that the initiation of reprogramming is not tightly coupled to OSK binding specificity. This result is supported by reports which show that substituting *OCT4* with mutant *OCT4* or orthologous *OCT6* in a reprogramming cocktail is able to initiate reprogramming in a manner similar to *OCT4*^52,53^.

### Affinity-dependent decay explains transient element kinetics

We show that the accessibility at transient elements decays in an affinity-dependent manner such that the sites with weaker motifs close earlier (**Fig 7ii**), suggesting that reduced expression of OSK at least in part explains the silencing of transient elements. Our motif analyses also suggest that stage-specific expression of additional TFs such as TFAP2C, ZIC, and GRHL determines subsequent chromatin dynamics (**Fig 7iv,v**). Such intricate and granular analyses of stage-specific sequence drivers of chromatin accessibility and the footprints they generate underscores the importance of base-resolution deep learning models in the study of cis-regulation.

### Evidence for a functional role of transient peaks in suppressing the fibroblast program via competitive redistribution of AP-1

Our study provides new insights into the role of Early Transient peaks in somatic silencing. Previous work has clearly demonstrated that fibroblast reprogramming is accelerated by inhibiting the expression or binding of somatic TFs belonging to the AP-1 family, and conversely arrested upon increasing AP-1 levels^54,55^. However, the mechanism underlying rapid somatic silencing in reprogramming remains nebulous.

The ChromBPNet models revealed that Early Transient peaks harbor thousands of instances of occupied AP-1 motifs, supported by discernable footprints (**Fig 6a, 7ii**). These findings are consistent with previous studies that have proposed a mechanism of “repression-by-theft”, wherein transient elements mediate the redistribution of somatic TFs away from their somatic targets^7,9^. This mechanism is analogous to the silencing of Runx1 targets caused by redistribution of Runx1 upon expression of the pioneer factor *SPI1* in T-cell development^5,56^, and the loss of pluripotency observed when introducing exogenous OCT4 binding arrays in ESCs^57^.

The single-nucleus multiome data at the earliest time points, revealed a quantitative association between the sequestration of somatic TFs to newly opened sites and the repression of somatic gene expression within individual nuclei. Our analysis provides the first direct, single-cell evidence supporting the “repression-by-theft” hypothesis. Our analyses suggest that this association is likely mediated by OSK stoichiometry, and may explain why sustained supraphysiological expression of the reprogramming factors is crucial for productive reprogramming. These findings not only enhance our understanding of reprogramming, but also have the potential to inform research on partial reprogramming, which involves the overexpression of reprogramming factors for a short interval of time^58^.

### A generalizable framework for deciphering regulation of natural and engineered cell fate

Together, this study provides a template for studying the effects of heterogeneous stoichiometry on cis-regulation. We exploit the stochasticity of Sendai virus induction to study the combinatorial state space of TF overexpression in reprogramming, and leverage powerful neural network models that learn cell state, TF stoichiometry-aware mappings from DNA sequence to chromatin accessibility. Our approach outlines a powerful paradigm for interrogating the stoichiometric and cis-regulatory sequence determinants of factor-induced or knockout de-differentiation and trans-differentiation trajectories.

### Limitations and future directions

Our analysis is subject to certain limitations. First, the cell state transition trajectories identified in our study represent our best guess estimate by combining multiple lines of evidence including pseudotime inference, Sendai virus dilution, and findings from previous studies.

Validating these trajectories would necessitate the use of lineage tracing methods. Second, the peak-gene linking is approximate, in part due to computational linking of scATAC-seq and scRNA-seq cells. This can impact the accuracy of the T2G networks. Third, ChromBPNet models explain chromatin dynamics that are predicted by local 2kb sequence windows. The models may be inaccurate when chromatin dynamics are determined by long-range interactions, and changes to methylation and histone modifications. For example, dynamic CTCF peaks that are closed in fibroblasts and open over the course of reprogramming are incorrectly predicted to be constitutively accessible by the models. To partially overcome this limitation, we restrict our analysis to peaks that are accessible in the respective cell types.

Improvements in experimental, computational and modeling approaches will yield increasingly accurate and precise insights into the molecular mechanisms underlying cellular reprogramming. That said, our analytic approach should prove of broad utility and pave the way for new analyses and insights into the role of TFs in directing cellular reprogramming that may have previously been missed.

## Acknowledgements

This work was supported by grants from the NIH R01HG009674 (A.K. and H.M.B.), R01GM136737 (K.C.W.) and R61AR076815 (K.C.W.). This work was supported by the Milky Way Research Foundation (A.K.). K.C.W. is a New York Stem Cell Foundation–Robertson Investigator, and the Stephen Bechtel Endowed Faculty Scholar in Pediatric Translational Medicine, Stanford Maternal and Child Health Research Institute. M.A. is supported by the NSF Graduate Research Fellowship Program.

All icons in schematics are obtained from BioRender.com. The computing for this project was performed in part on the Sherlock cluster. We thank Stanford University, and the Stanford Research Computing Center for computational resources.

## Author contributions

S.N., M.A., I.K., K.C.W., and A.K. conceived the project and designed experiments. M.A. performed experiments. S.N. performed the analyses with inputs from A.K., L.S., A.P., J.S. A.B., Y.X.W., D.B., H.B., and K.C.W. S.N. and A.K. wrote the manuscript.

## Data, resource and code availability

- Aligned fragment files for scATAC-seq and multiome, and unfiltered counts matrices for scRNA-seq and multiome are deposited in the Gene Expression Omnibus database with the Super-Series reference number <u>GSE242424</u>. The mapped reads are also available at https://doi.org/10.5281/zenodo.8294147. Note that raw reads are not available due to patient privacy concerns.

- Analysis products: https://doi.org/10.5281/zenodo.8313961 (includes counts matrices for scATAC-seq, scRNA-seq and multiome integrated across all samples, cell representations, UMAP coordinates, cluster assignment. For scATAC-seq clusters, includes per cluster fragment files and peak calls. For scRNA-seq, contains endogenous and Sendai OSKM estimates.)

- ChromBPNet models and data: https://doi.org/10.5281/zenodo.8299710

- Counts and profile importance scores, corresponding TF-MoDISco outputs, and consolidated motifs: https://doi.org/10.7303/syn52331899

- Interactive resources: https://kundajelab.github.io/reprogramming-browser/home.html contains interactive resources including a scRNA-seq browser and scATAC-seq cluster browser.

- Code for analyses and model training, evaluation and interpretation: https://github.com/kundajelab/scATAC-reprog

## Supplementary table captions

**Supp Table 1**: Peak-gene links with FDR < 1e-4. For analyses in the paper, we filter to peaks with absolute correlation >0.45.

**Supp Table 2**: Gene sets and GO enrichment results from gProfiler2 for each peak set (**Methods**). For each peak set, genes that are linked to peaks are ranked based on the number of peaks with which they are linked. gProfiler2 terms with source “TF” are excluded.

## Software and resources used

- ENCODE ATAC-seq and Chip-seq pipelines^59^

- ArchR^60^

- 10x Genomics Cell Ranger 6.0.2^61^

- GViz^62^

- BentoBox (now PlotGardener)^63^

- SnapATAC^64^

- Chromap^65^

- ggseqlogo^66^

- logomaker^67^

- gProfiler2^68^

- cistromeDB^69^

- Seurat^70^

- DoubletFinder^71^

- PAGA^34^

- Scanpy^72^

- Harmony^73^

- AMULET^74^

- ChromVAR^36^

- ClusterR^75^

- DESeq2^76^

- GimmeMotifs^77^

- Tomtom^78^

- HOMER^79^

