## Supplementary Figures for "Deep learning the cis-regulatory code of chromatin dynamics during cellular reprogramming"

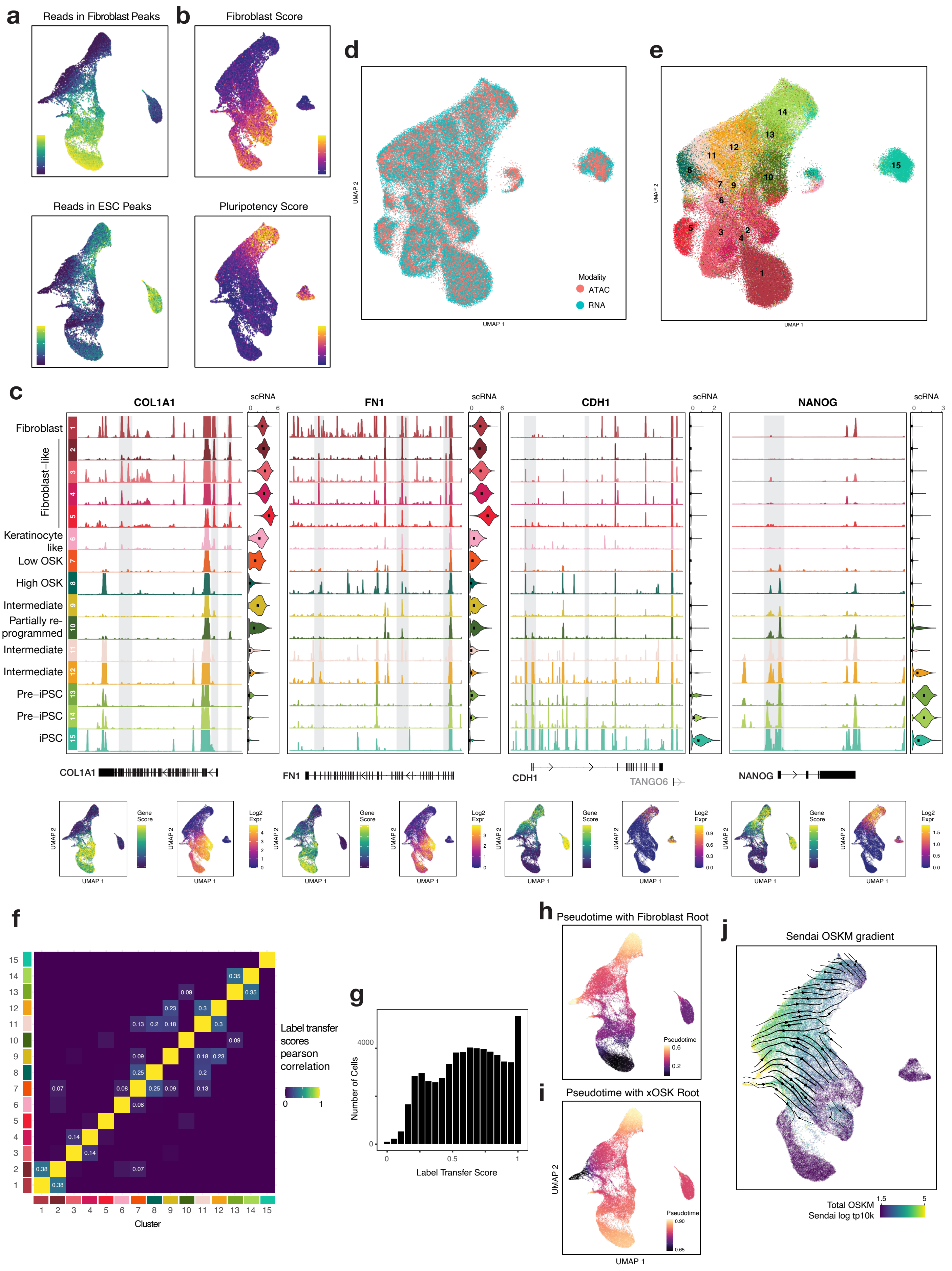
a
Reads in Fibroblast Peaks
Reads in ESC Peaks
b
Fibroblast Score
Pluripotency Score
d
Modality
ATAC
RNA
UMAP 1
UMAP 2
e
c
COL1A1
FN1
CDH1
NANOG
scRNA
Fibroblast
Fibroblast-like
Keratinocyte like
Low OSK
High OSK
Intermediate
Partially re-programmed
Intermediate
Intermediate
Pre–iPSC
Pre–iPSC
iPSC
TANGO6
Gene Score
Log2 Expr
f
Label transfer scores pearson correlation
Cluster
g
Number of Cells
Label Transfer Score
h
Pseudotime with Fibroblast Root
Pseudotime
i
Pseudotime with xOSK Root
j
Sendai OSKM gradient
Total OSKM Sendai log tp10k

**Supp Fig 1:**
**a)** Per-cell score based on overlap of reads with all fibroblast (top) or Primed ESC (bottom) peaks **b)** Per-cell averaged gene-expression z-scores based on predefined sets of 84 fibroblast-specific (top) and 20 iPSC-specific (bottom) genes **c)** Locus plots for COL1A1, FN1, CDF1, and NANOG, along with their scATAC-seq derived gene scores and scRNA-seq gene expression **d)** Joint CCA embedding of scRNA-seq and scATAC-seq cells labeled by modality **e)** Same as d), labeled by cell state **f)** Correlation between cell state label transfer scores across scRNA-seq cells **g)** Histogram of cell state label transfer score assigned to the highest cell state for each scRNA-seq cell **h)** Pseudotime values with a fibroblast cell set as root **i)** Pseudotime values with Day 2 xOSK cell set as root **j)** This plot shows the direction of decreasing total OSKM Sendai expression over the scRNA-seq UMAP using arrows, excluding over cells with low Sendai expression. Under the assumption that Sendai expression decreases over time, these lines impose constraints over possible cell trajectories, such that cells may move perpendicular to the flow lines, but not against them.

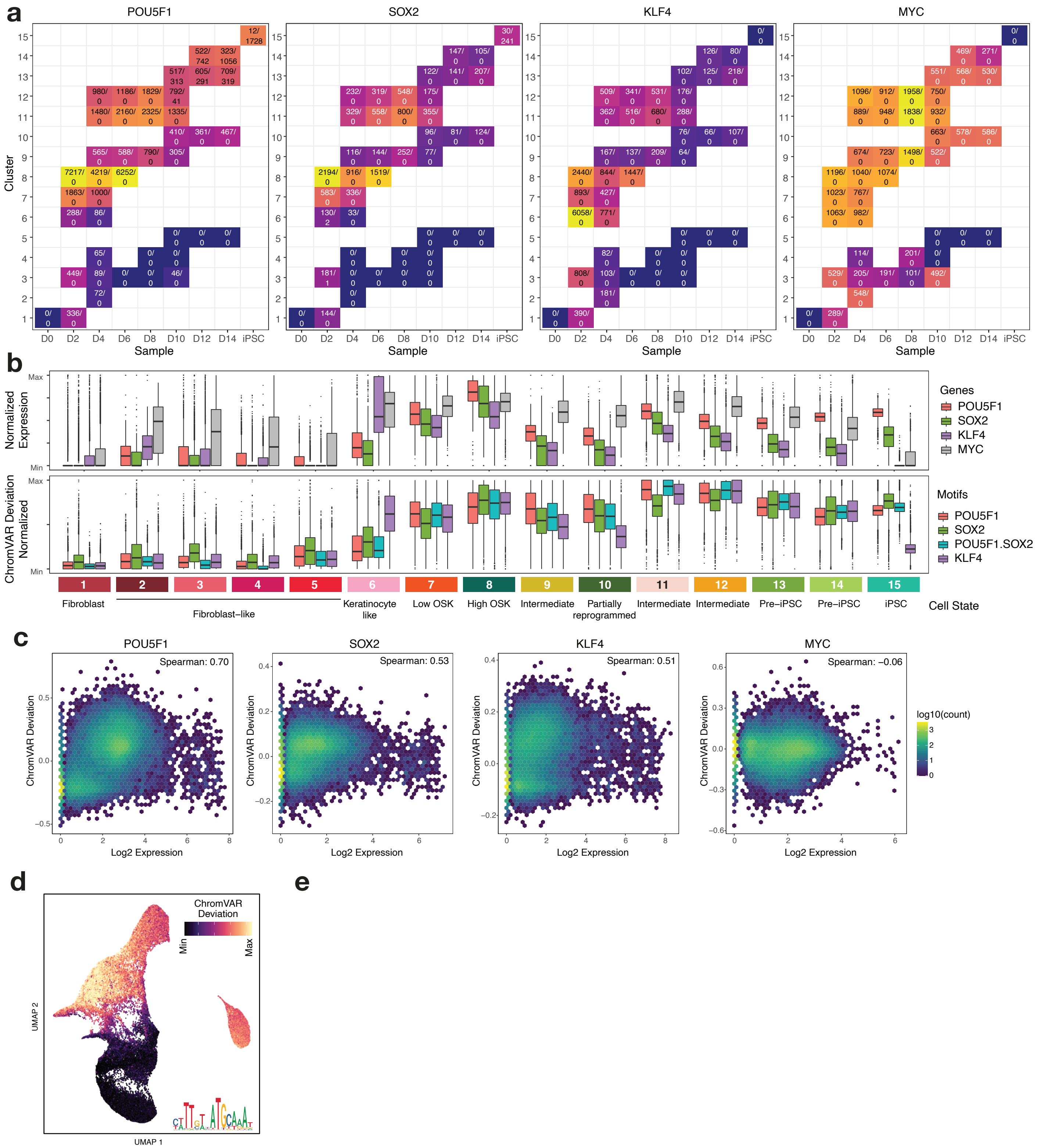
a
POU5F1
SOX2
KLF4
MYC
Cluster
Sample
D0 D2 D4 D6 D8 D10 D12 D14 iPSC
b
Normalized Expression
Max
Min
Genes
POU5F1
SOX2
KLF4
MYC
ChromVAR Deviation Normalized
Motifs
POU5F1
SOX2
POU5F1.SOX2
KLF4
Fibroblast
Fibroblast–like
Keratinocyte like
Low OSK
High OSK
Intermediate
Partially reprogrammed
Intermediate
Intermediate
Pre–iPSC
Pre–iPSC
iPSC
Cell State
c
POU5F1
Spearman: 0.70
SOX2
Spearman: 0.53
KLF4
Spearman: 0.51
MYC
Spearman: –0.06
ChromVAR Deviation
Log2 Expression
log10(count)
d
ChromVAR Deviation
Min
Max
UMAP 2
UMAP 1
e

**Supp Fig 2:**
**a)** Median estimated Sendai and endogenous expression of OSKM (in TPM) of cells stratified by time point and cell state. Only configurations with >50 cells are shown. Within each cell, the first value refers to the Sendai median expression and the second value refers to the endogenous median expression. Sendai is set as median total - median endogenous **b)** Box plots of per state normalized gene expression of OSKM (top) and ChromVAR deviation for POU5F1, SOX2, POU5F1-SOX2 and KLF4 motifs (bottom). For genes, the expression is min-max normalized and outliers outside the 99th percentile are clipped. For motifs, the ChromVAR deviations are min-max normalized and outliers below the 10th percentile and above the 95th percentile are clipped **c)** Density plots of gene expression of OSKM versus ChromVAR deviations for their respective motifs across matched ATAC-RNA cells **d)** ChromVAR deviation scores for the OCT-SOX motif

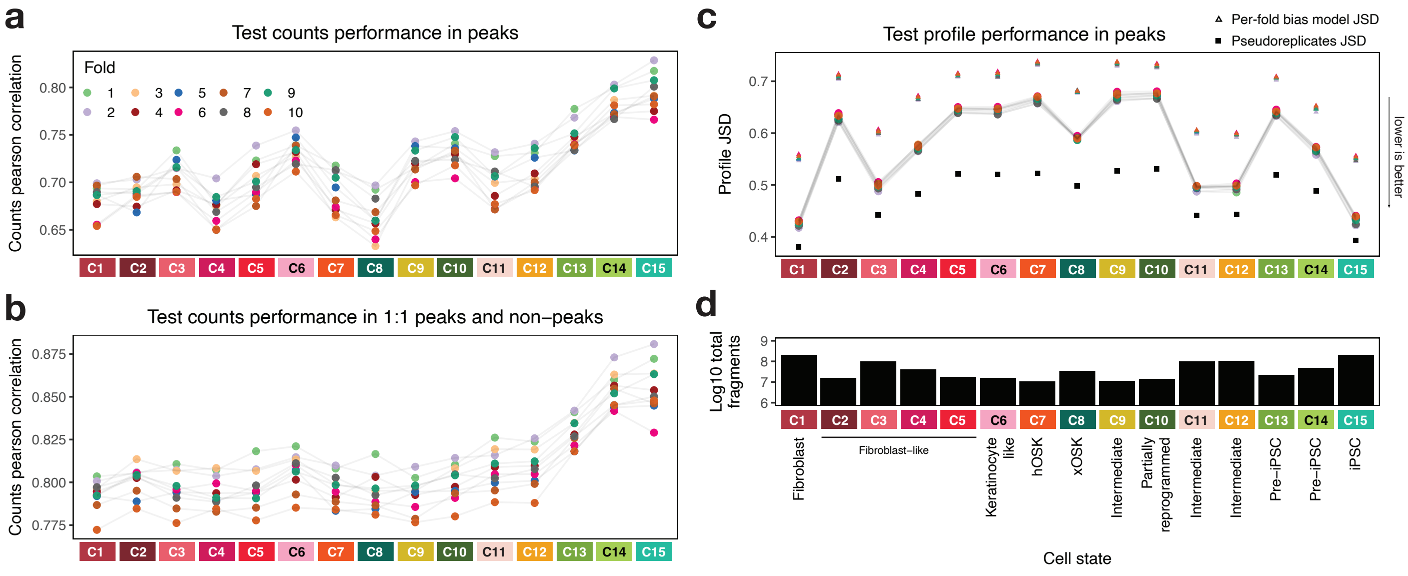


**Supp Fig 3:**
**a)** ChromBPNet counts predictions Pearson correlation with observed reads in test peak regions for all 10 model folds. Note that peaks are cell state specific **b)** ChromBPNet counts predictions Pearson correlation with observed reads in test peak and GC-matched non-peak regions (equal number of peaks and non-peaks) for all 10 model folds **c)** ChromBPNet profile prediction performance of the models showing the median JSD (lower is better) between predicted and observed base-resolution Tn5 insertion distributions across peaks. Triangles show the prediction of the bias model on each fold's test set, which serves as a worst-case bound. Note that the bias model is fixed but the test regions are different for each fold. The black squares show the JSD between 2 independently drawn samples of half the read depth for each sample, and serves as a reasonable best-case bound. Note that it is difficult to compare profile performance across cell states as the JSD depends on read depth of the samples, which determines the ground truth **d)** Log10 fragment counts of each sample, which correlates with sample-specific JSD in c)

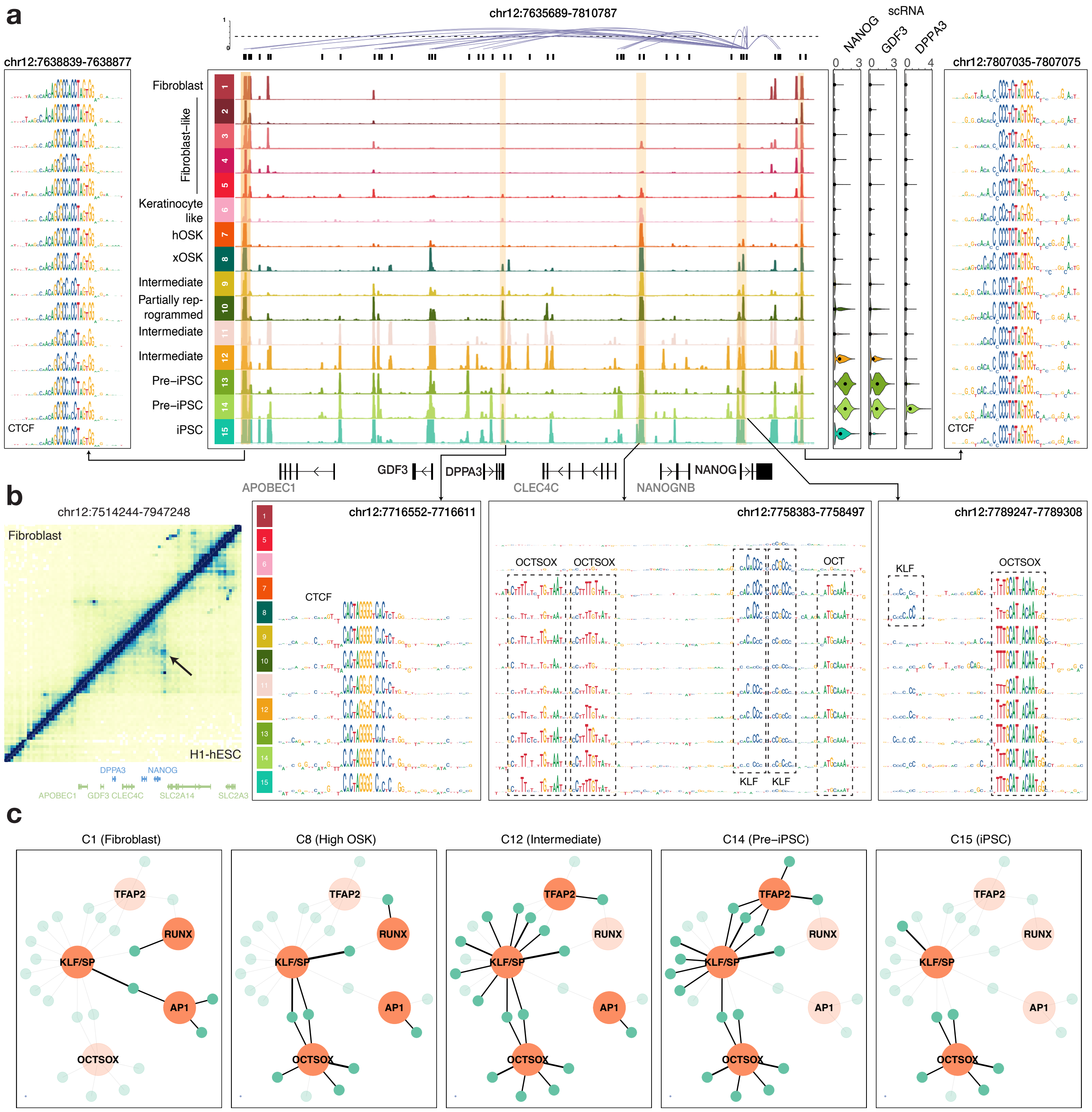


**Supp Fig 4:**

**a)** Extended version of **Fig 3d**. Contribution scores are shown only for cell states for which the locus was called as a peak **b)** Micro-C contact maps in human foreskin fibroblasts (4DNFIPC7P27B) and H1-hESCs (4DNFI2TK7L2F) from (Krietenstein et al. 2020). The arrow highlights a loop found in H1-hESC but not fibroblasts **c)** Extended version of **Fig 3e**

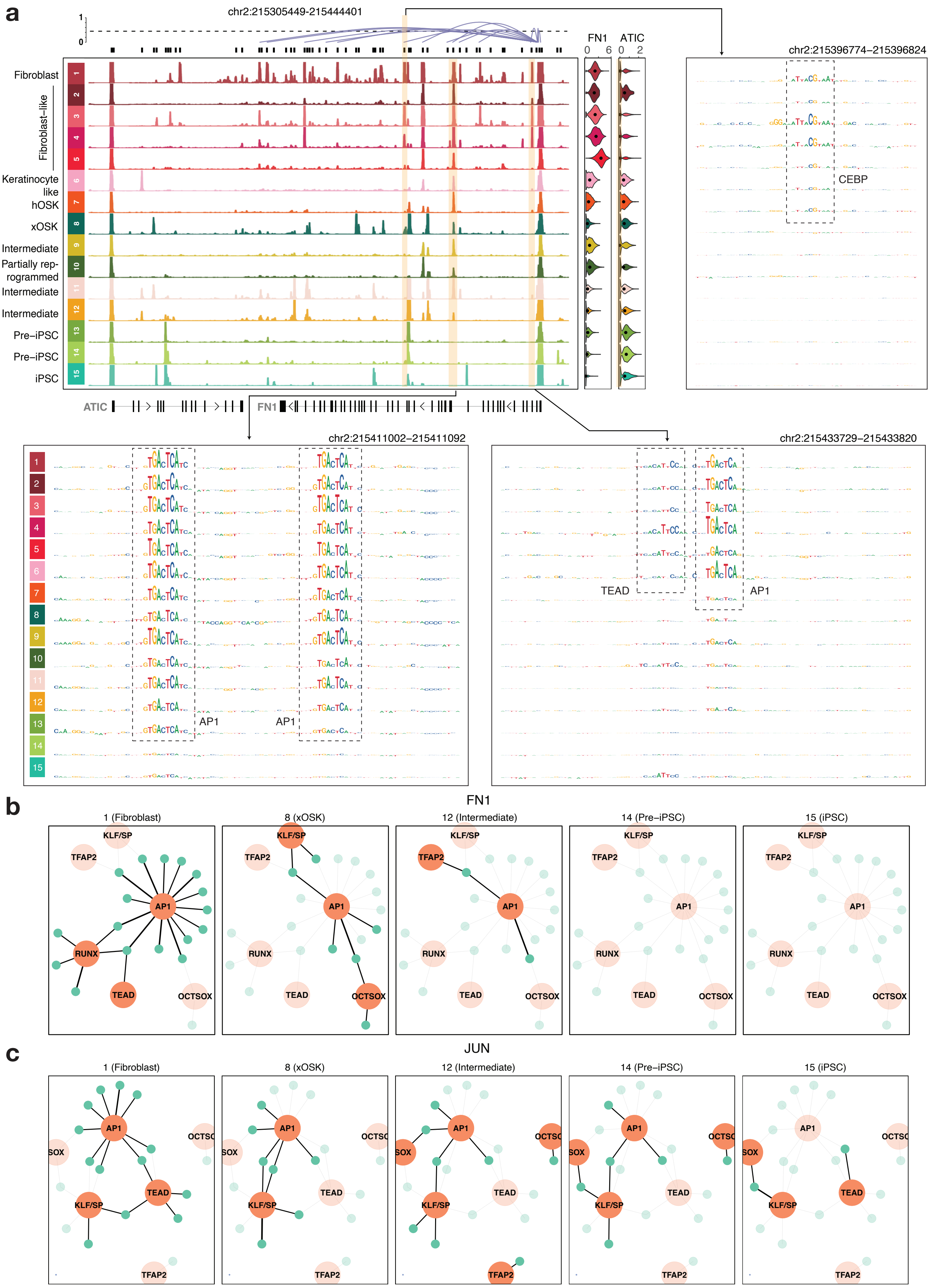

a
chr2:215305449-215444401
1
0
FN1
ATIC
0
6
0
2
chr2:215396774–215396824
Fibroblast
Fibroblast–like
Keratinocyte like
hOSK
xOSK
Intermediate
Partially rep-rogrammed
Intermediate
Intermediate
Pre–iPSC
Pre–iPSC
iPSC
CEBP
ATIC
FN1
chr2:215411002–215411092
chr2:215433729–215433820
TEAD
AP1
AP1
AP1
b
FN1
1 (Fibroblast)
8 (xOSK)
12 (Intermediate)
14 (Pre–iPSC)
15 (iPSC)
KLF/SP
TFAP2
AP1
RUNX
TEAD
OCTSOX
c
JUN
1 (Fibroblast)
8 (xOSK)
12 (Intermediate)
14 (Pre–iPSC)
15 (iPSC)
AP1
OCTSO
SOX
TEAD
KLF/SP
TFAP2

**Supp Fig 5:**
**a)** Locus plot for the FN1 gene (similar to **Fig 3d**) Contribution scores are shown for all cell states **b)** TF-to-gene network of FN1 (similar to **Fig 3e**) **c)** TF-to-gene network of JUN

a

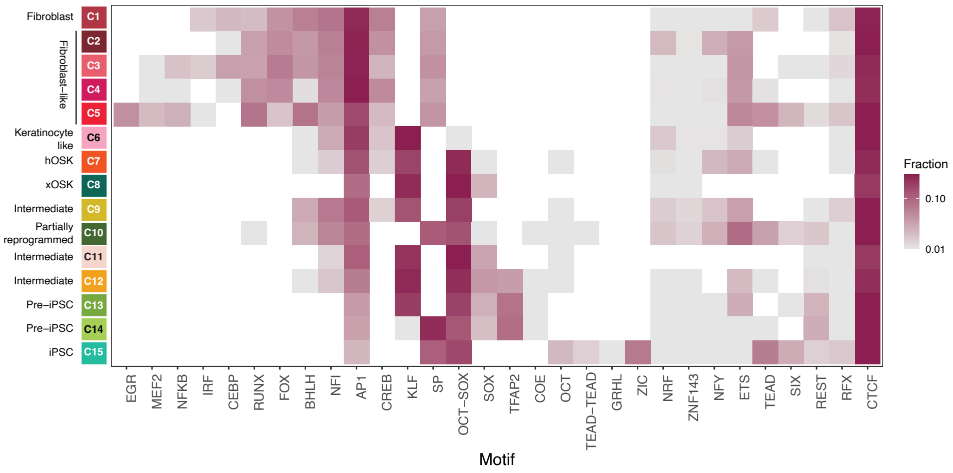


**Supp Fig 6:**

**a)** Consolidated motifs extracted by TF-MoDISco from each cell state. While each cell state cluster extracts slightly different motifs, these were grouped and consolidated into a set of 30 non-redundant motifs. Each tile represents the log10 fraction of number of instances extracted by TF-MoDISco for that motif across instances of all motifs for that cell state. Empty (white) tiles indicate that the motif was not detected in that cell state (with at least 50 instances). Note that KLF and SP are very similar motifs, and the motif assignments to them are nearly exclusive based on which one was a closer match.

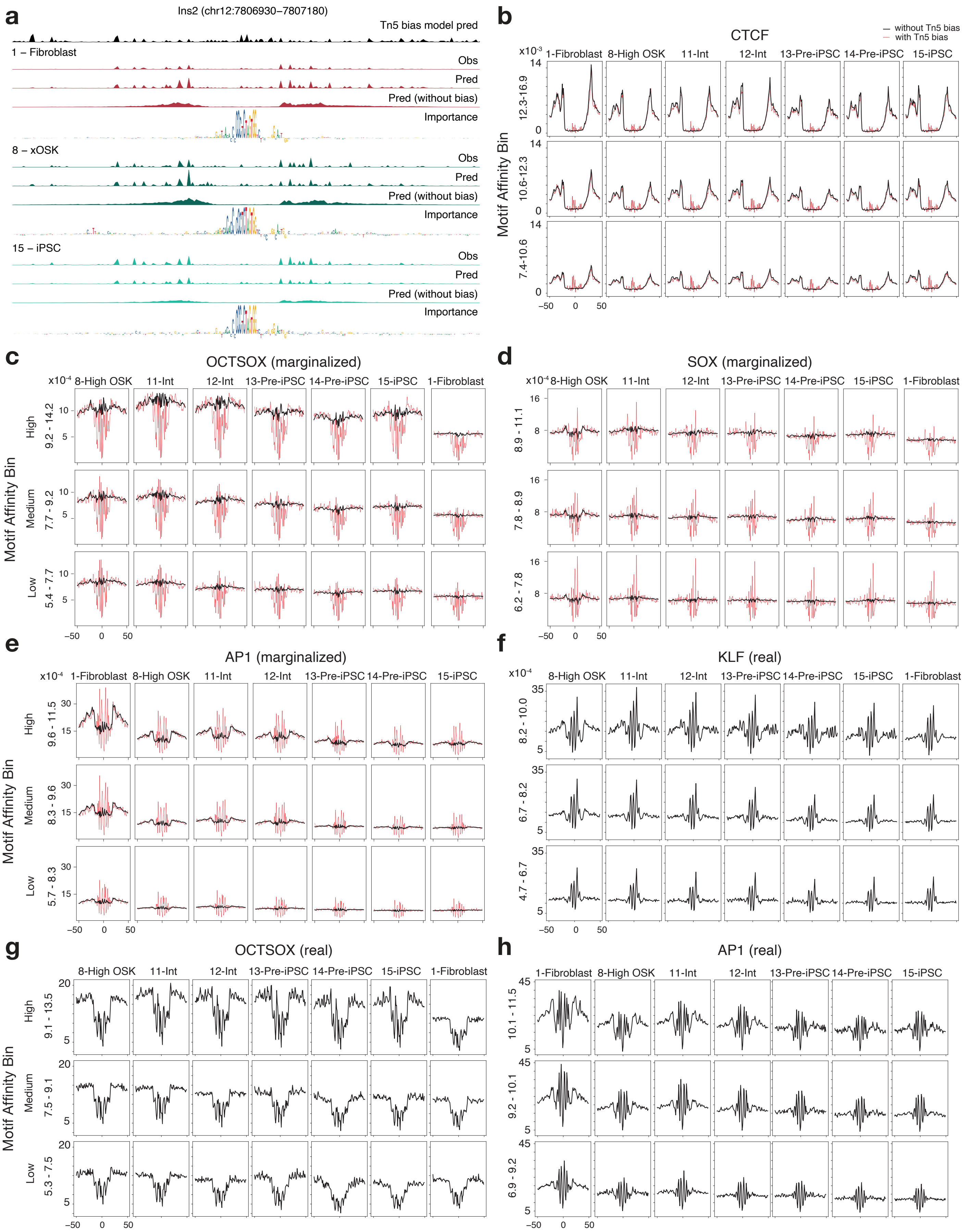
a
Ins2 (chr12:7806930–7807180)
Tn5 bias model pred
1 – Fibroblast
Obs
Pred
Pred (without bias)
Importance
8 – xOSK
15 – iPSC
b
CTCF
without Tn5 bias
with Tn5 bias
1-Fibroblast 8-High OSK 11-Int 12-Int 13-Pre-iPSC 14-Pre-iPSC 15-iPSC
Motif Affinity Bin
12.3-16.9
10.6-12.3
7.4-10.6
c
OCTSOX (marginalized)
d
SOX (marginalized)
e
AP1 (marginalized)
f
KLF (real)
g
OCTSOX (real)
h
AP1 (real)
High
Medium
Low

**Supp Fig 7:**
**a)** Zoom in of Ins2 locus from **Fig 3d**, with the bias model prediction at the top and normalized observed and predicted (with and without bias) profiles, and the counts contribution track for 3 states **b)** Virtual footprinting for CTCF obtained by inserting instances of CTCF stratified by log-odds scores into random background sequences and averaging predicted profile probability distributions with (red) and without (black) bias (**Methods**) for each cell state **c)** Same as b) for OCT-SOX motif. States ordered in approximately decreasing order of OCT4 and SOX2 concentration **d)** Same as c) for SOX motif **e)** Same as c) for AP1 motif **f)** Footprints derived by aggregating normalized observed reads over motif instances of KLF stratified by log-odds scores for each cell state. States ordered in decreasing order of KLF4 concentration **g)** Same as f) for OCT-SOX motif **h)** Same as f) for AP1 motif

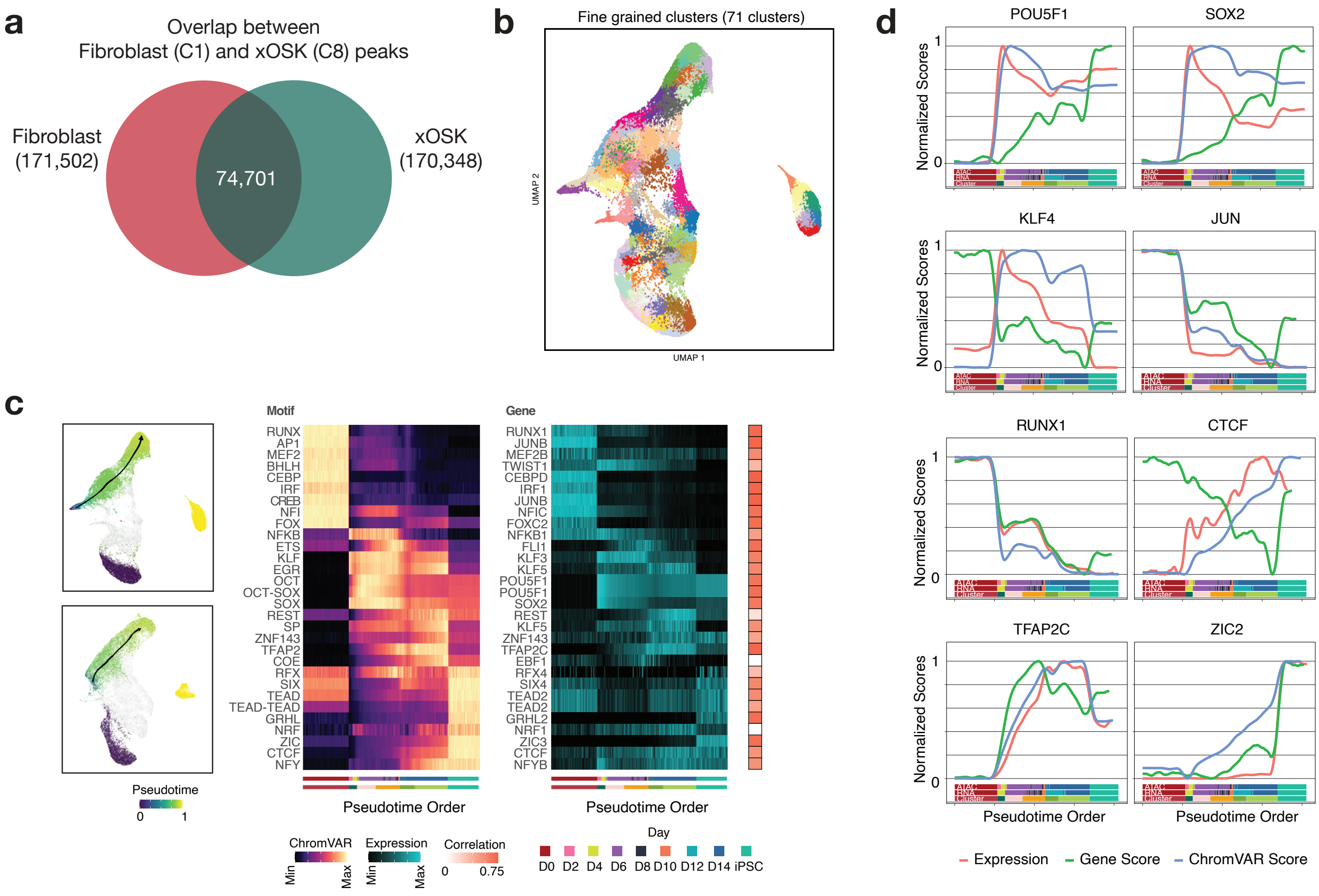


**Supp Fig 8:**

**a)** Venn diagram of Fibroblast (C1) and xOSK (C8) peaks **b)** 71 clusters of cells representing higher resolution subpopulations **c)** ChromVAR deviation scores for all motifs across the primary reprogramming trajectory, along with gene expression of the gene within the motif family with the highest correlation with ChromVAR deviation score. Colored strips below heatmaps show day and cell state labels along the pseudotime trajectory. Red tiles show correlation between ChromVAR deviation and gene expression. **d)** Normalized gene expression, chromatin-derived gene score, and ChromVAR deviation score for selected TFs along primary reprogramming trajectory

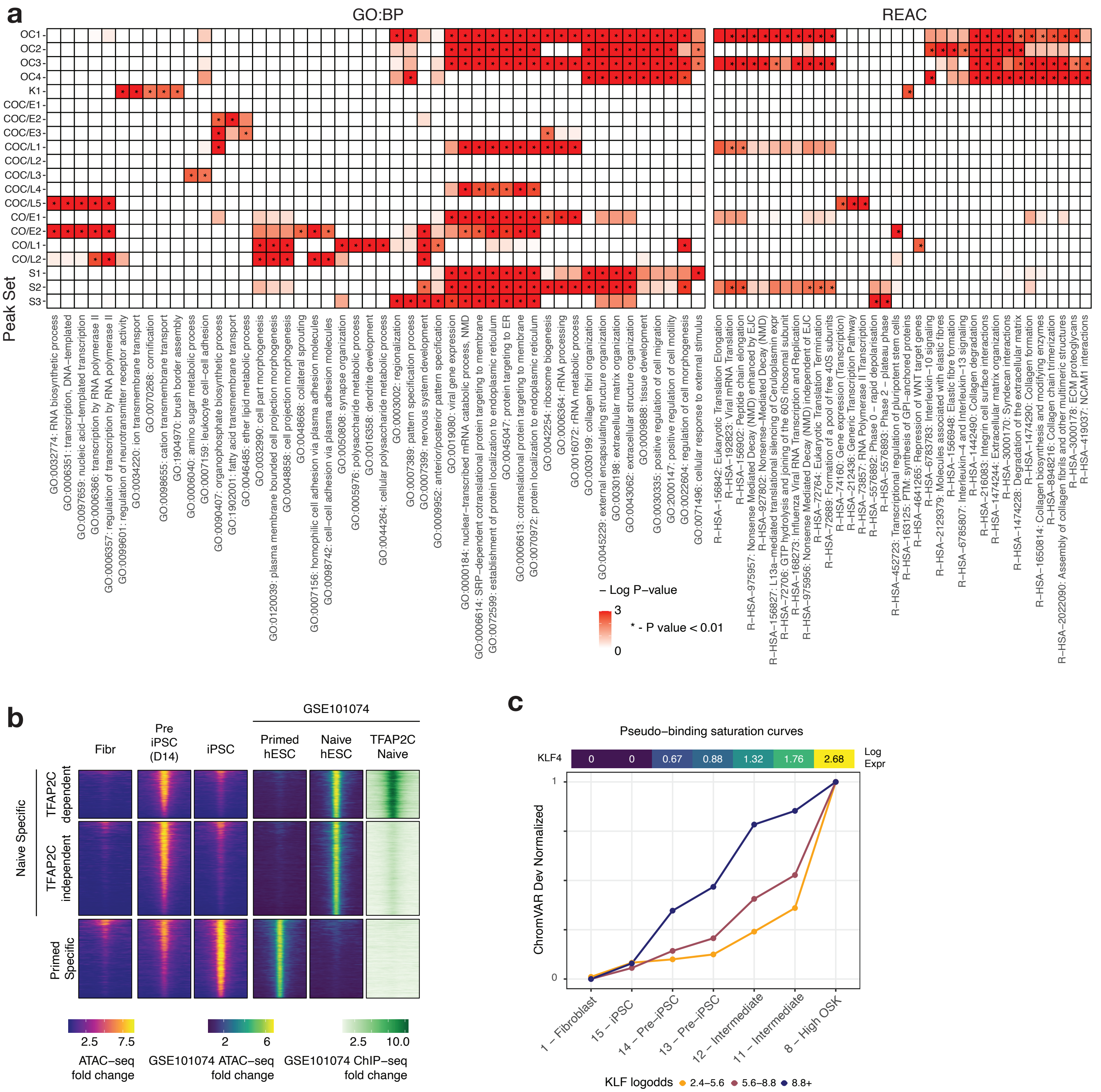

**Supp Fig 9:**
**a)** Curated GO and REAC terms enriched across peak sets **b)** Integration of scATAC-seq data with naive and primed hESC ATAC-seq and naive hESC TFAP2C Chip-seq data (GSE101074) **c)** Same as **Fig 5f** for KLF4

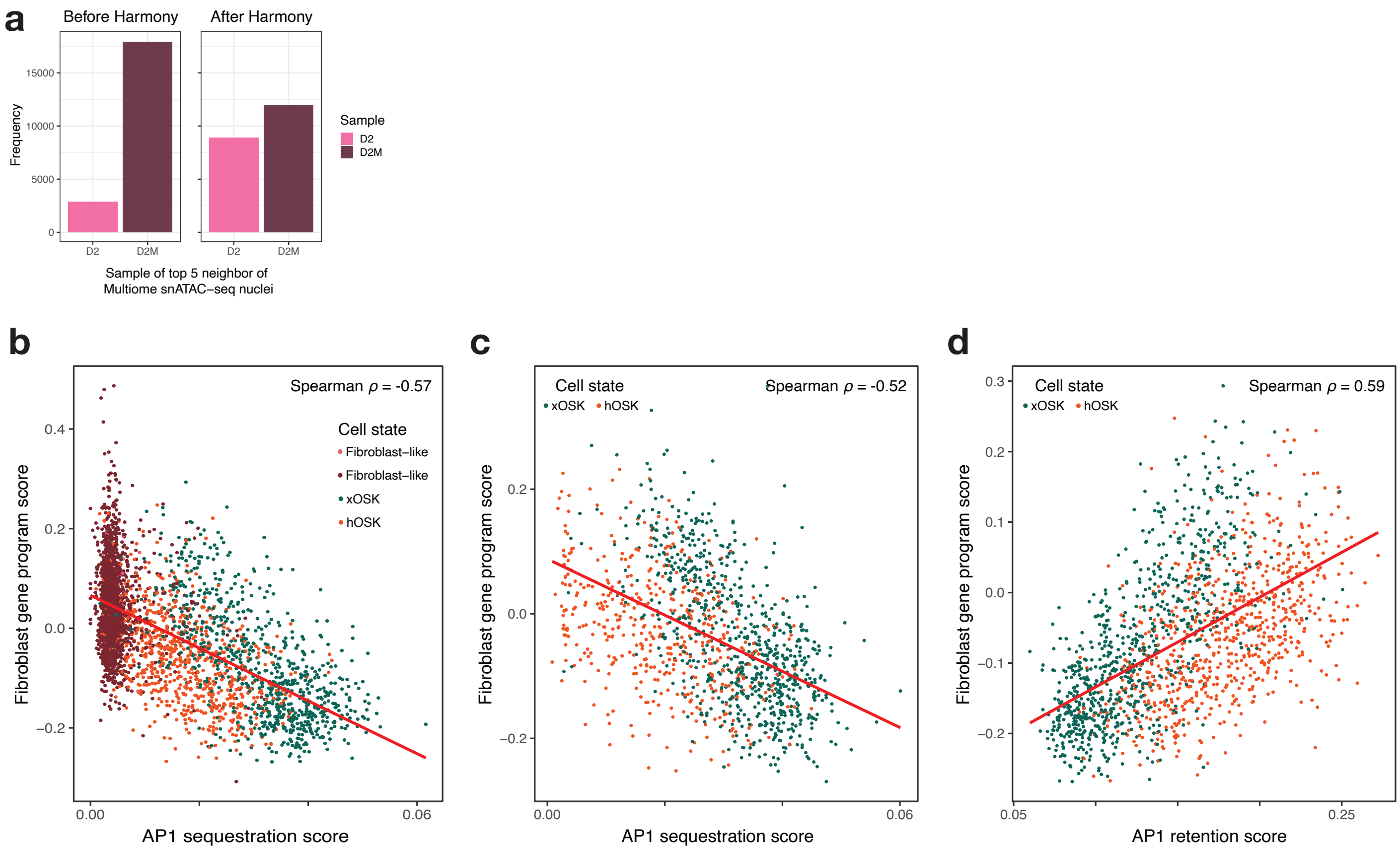


**Supp Fig 10:**

**a)** Sample identity (scATAC-seq D2 or sn-multiome D2M) of top 5 neighbors of Multiome snATAC-seq nuclei before and after Harmony correction for technology **b)** Similar to **Fig 6g**, but points are subsetted to Day 2 multiome nuclei from fibroblast-like, hOSK and xOSK states (Spearman $p$-value $< 2.2 \times 10^{-16}$) **c)** Similar to **Fig 6g**, but points are subsetted to Day 1 multiome nuclei from hOSK and xOSK states (Spearman $p$-value $< 2.2 \times 10^{-16}$) **d)** Scatter-plot of AP-1 retention score versus snRNA-seq expression of fibroblast-specific genes. Each point is a single nucleus from Day 2 multiome data from hOSK and xOSK states (Spearman $p$-value $< 2.2 \times 10^{-16}$)
