## Extended Figures for "Deep learning the cis-regulatory code of chromatin dynamics during cellular reprogramming"

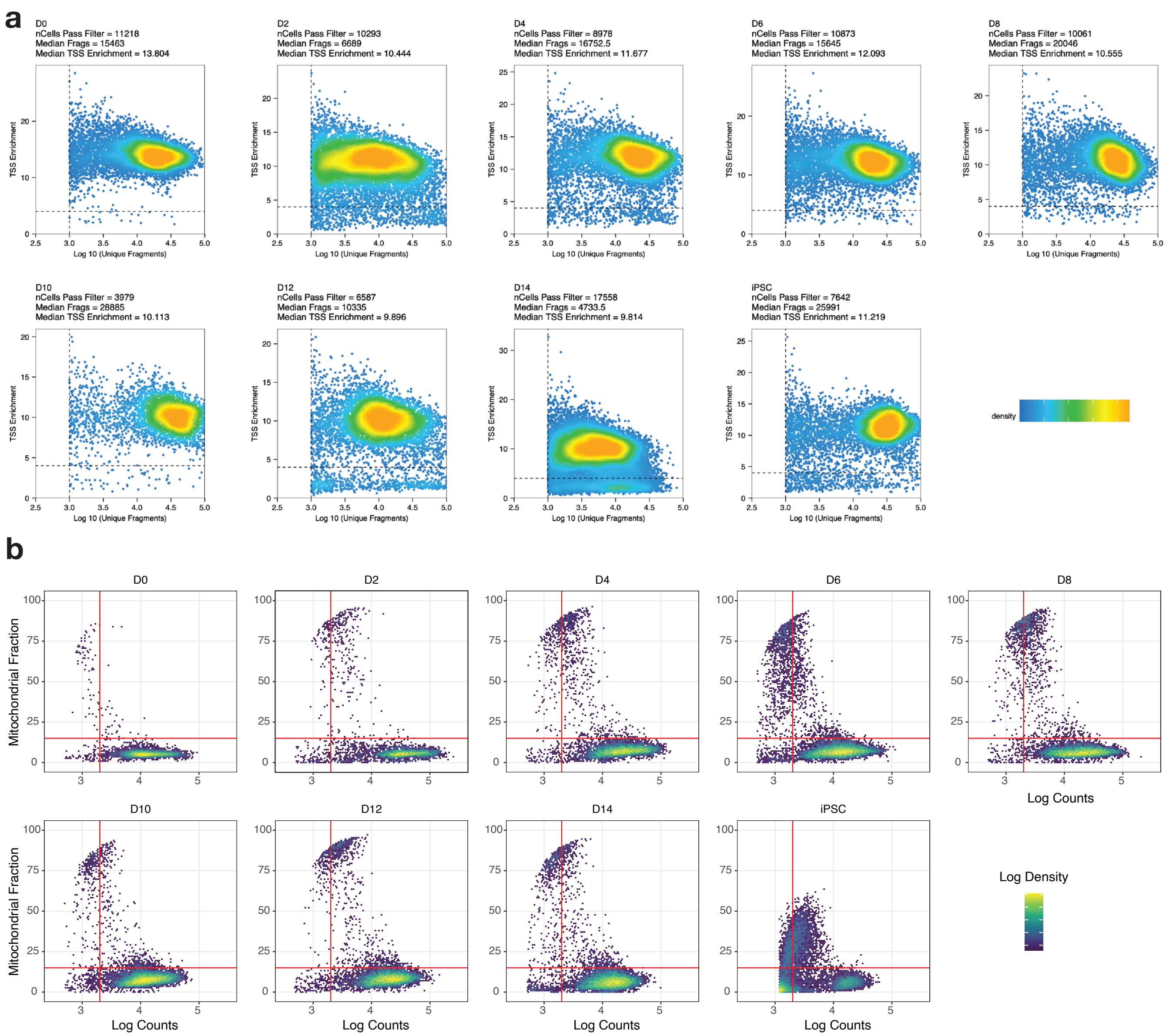


**Extended Fig 1:**
**a)** Quality control for scATAC-seq samples **b)** Quality control for scRNA-seq samples

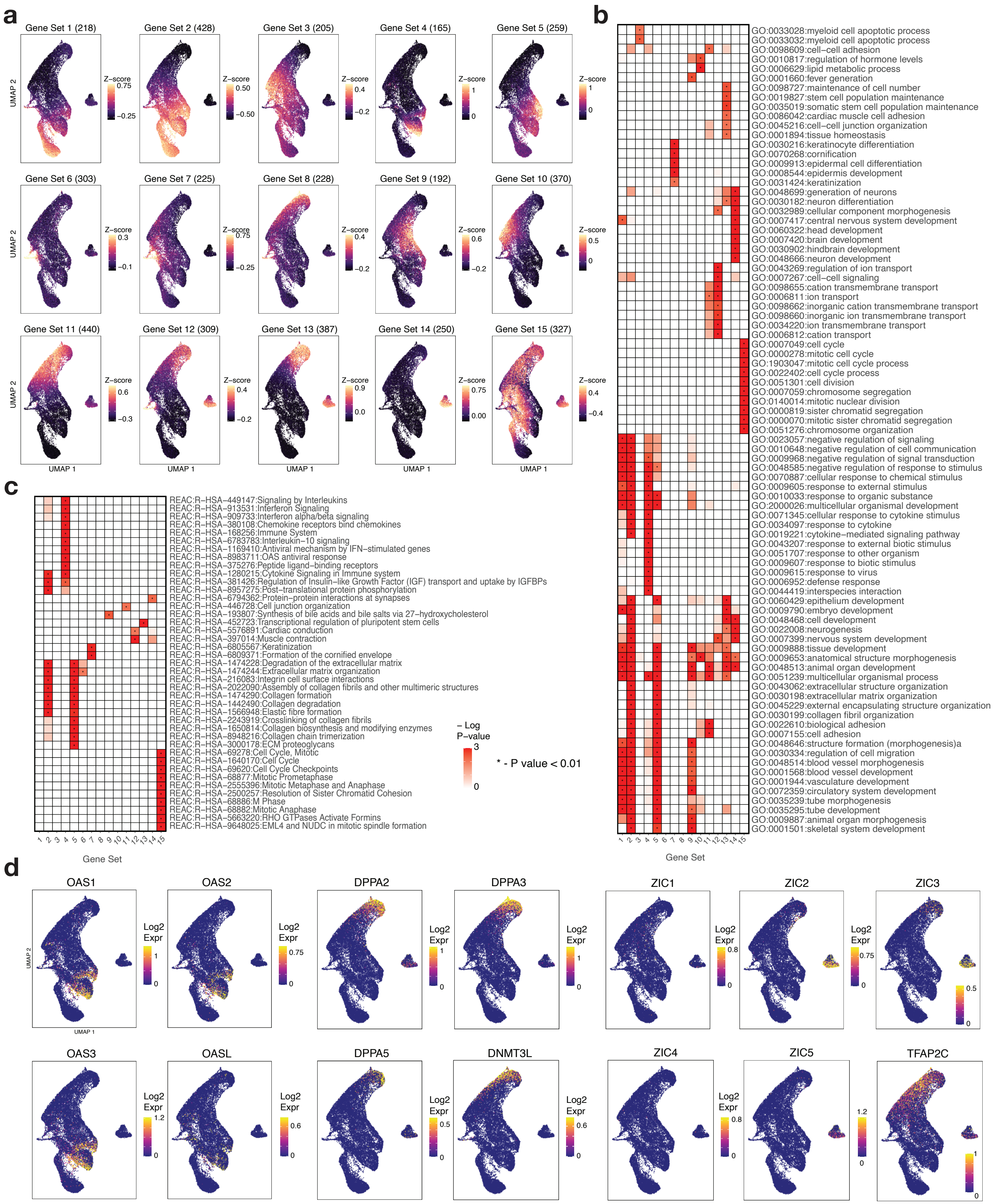

a
Gene Set 1 (218)
Gene Set 2 (428)
Gene Set 3 (205)
Gene Set 4 (165)
Gene Set 5 (259)
Gene Set 6 (303)
Gene Set 7 (225)
Gene Set 8 (228)
Gene Set 9 (192)
Gene Set 10 (370)
Gene Set 11 (440)
Gene Set 12 (309)
Gene Set 13 (387)
Gene Set 14 (250)
Gene Set 15 (327)
Z−score
UMAP 2
UMAP 1
b
GO:0033028:myeloid cell apoptotic process
GO:0033032:myeloid cell apoptotic process
GO:0098609:cell−cell adhesion
GO:0010817:regulation of hormone levels
GO:0006629:lipid metabolic process
GO:0001660:fever generation
GO:0098727:maintenance of cell number
GO:0019827:stem cell population maintenance
GO:0035019:somatic stem cell population maintenance
GO:0086042:cardiac muscle cell adhesion
GO:0045216:cell−cell junction organization
GO:0001894:tissue homeostasis
GO:0030216:keratinocyte differentiation
GO:0070268:cornification
GO:0009913:epidermal cell differentiation
GO:0008544:epidermis development
GO:0031424:keratinization
GO:0048699:generation of neurons
GO:0030182:neuron differentiation
GO:0032989:cellular component morphogenesis
GO:0007417:central nervous system development
GO:0060322:head development
GO:0007420:brain development
GO:0030902:hindbrain development
GO:0048666:neuron development
GO:0043269:regulation of ion transport
GO:0007267:cell−cell signaling
GO:0098655:cation transmembrane transport
GO:0006811:ion transport
GO:0098662:inorganic cation transmembrane transport
GO:0098660:inorganic ion transmembrane transport
GO:0034220:ion transmembrane transport
GO:0006812:cation transport
GO:0007049:cell cycle
GO:0000278:mitotic cell cycle
GO:1903047:mitotic cell cycle process
GO:0022402:cell cycle process
GO:0051301:cell division
GO:0007059:chromosome segregation
GO:0140014:mitotic nuclear division
GO:0000819:sister chromatid segregation
GO:0000070:mitotic sister chromatid segregation
GO:0051276:chromosome organization
GO:0023057:negative regulation of signaling
GO:0010648:negative regulation of cell communication
GO:0009968:negative regulation of signal transduction
GO:0048585:negative regulation of response to stimulus
GO:0070887:cellular response to chemical stimulus
GO:0009605:response to external stimulus
GO:0010033:response to organic substance
GO:2000026:multicellular organismal development
GO:0071345:cellular response to cytokine stimulus
GO:0034097:response to cytokine
GO:0019221:cytokine−mediated signaling pathway
GO:0043207:response to external biotic stimulus
GO:0051707:response to other organism
GO:0009607:response to biotic stimulus
GO:0009615:response to virus
GO:0006952:defense response
GO:0044419:interspecies interaction
GO:0060429:epithelium development
GO:0009790:embryo development
GO:0048468:cell development
GO:0022008:neurogenesis
GO:0007399:nervous system development
GO:0009888:tissue development
GO:0009653:anatomical structure morphogenesis
GO:0048513:animal organ development
GO:0051239:multicellular organismal process
GO:0043062:extracellular structure organization
GO:0030198:extracellular matrix organization
GO:0045229:external encapsulating structure organization
GO:0030199:collagen fibril organization
GO:0022610:biological adhesion
GO:0007155:cell adhesion
GO:0048646:structure formation (morphogenesis)a
GO:0030334:regulation of cell migration
GO:0048514:blood vessel morphogenesis
GO:0001568:blood vessel development
GO:0001944:vasculature development
GO:0072359:circulatory system development
GO:0035239:tube morphogenesis
GO:0035295:tube development
GO:0009887:animal organ morphogenesis
GO:0001501:skeletal system development
Gene Set
c
REAC:R−HSA−449147:Signaling by Interleukins
REAC:R−HSA−913531:Interferon Signaling
REAC:R−HSA−909733:Interferon alpha/beta signaling
REAC:R−HSA−380108:Chemokine receptors bind chemokines
REAC:R−HSA−168256:Immune System
REAC:R−HSA−6783783:Interleukin−10 signaling
REAC:R−HSA−1169410:Antiviral mechanism by IFN−stimulated genes
REAC:R−HSA−8983711:OAS antiviral response
REAC:R−HSA−375276:Peptide ligand−binding receptors
REAC:R−HSA−1280215:Cytokine Signaling in Immune system
REAC:R−HSA−381426:Regulation of Insulin−like Growth Factor (IGF) transport and uptake by IGFBPs
REAC:R−HSA−8957275:Post−translational protein phosphorylation
REAC:R−HSA−6794362:Protein−protein interactions at synapses
REAC:R−HSA−446728:Cell junction organization
REAC:R−HSA−193807:Synthesis of bile acids and bile salts via 27−hydroxycholesterol
REAC:R−HSA−452723:Transcriptional regulation of pluripotent stem cells
REAC:R−HSA−5576891:Cardiac conduction
REAC:R−HSA−397014:Muscle contraction
REAC:R−HSA−6805567:Keratinization
REAC:R−HSA−6809371:Formation of the cornified envelope
REAC:R−HSA−1474228:Degradation of the extracellular matrix
REAC:R−HSA−1474244:Extracellular matrix organization
REAC:R−HSA−216083:Integrin cell surface interactions
REAC:R−HSA−2022090:Assembly of collagen fibrils and other multimeric structures
REAC:R−HSA−1474290:Collagen formation
REAC:R−HSA−1442490:Collagen degradation
REAC:R−HSA−1566948:Elastic fibre formation
REAC:R−HSA−2243919:Crosslinking of collagen fibrils
REAC:R−HSA−1650814:Collagen biosynthesis and modifying enzymes
REAC:R−HSA−8948216:Collagen chain trimerization
REAC:R−HSA−3000178:ECM proteoglycans
REAC:R−HSA−69278:Cell Cycle, Mitotic
REAC:R−HSA−1640170:Cell Cycle
REAC:R−HSA−69620:Cell Cycle Checkpoints
REAC:R−HSA−68877:Mitotic Prometaphase
REAC:R−HSA−2555396:Mitotic Metaphase and Anaphase
REAC:R−HSA−2500257:Resolution of Sister Chromatid Cohesion
REAC:R−HSA−68886:M Phase
REAC:R−HSA−68882:Mitotic Anaphase
REAC:R−HSA−5663220:RHO GTPases Activate Formins
REAC:R−HSA−9648025:EML4 and NUDC in mitotic spindle formation
− Log P−value
* - P value < 0.01
Gene Set
d
OAS1
OAS2
DPPA2
DPPA3
ZIC1
ZIC2
ZIC3
OAS3
OASL
DPPA5
DNMT3L
ZIC4
ZIC5
TFAP2C
Log2 Expr

**Extended Fig 2:**
**a)** Gene expression Z-scores for 15 gene sets based on clustering of scRNA-seq data. Number of genes in each set indicated in parenthesis. These gene sets are distinct from the gene sets obtained by finding genes linked to peaks within the 20 peak sets **b)** Gene Ontology Biological Process terms enrichment matrix across the 15 gene sets **c)** Reactome (REAC) terms enrichment matrix across the 15 gene sets **d)** scRNA-seq gene expression plots for genes from the OAS, DPPA, and ZIC families and TFAP2C.

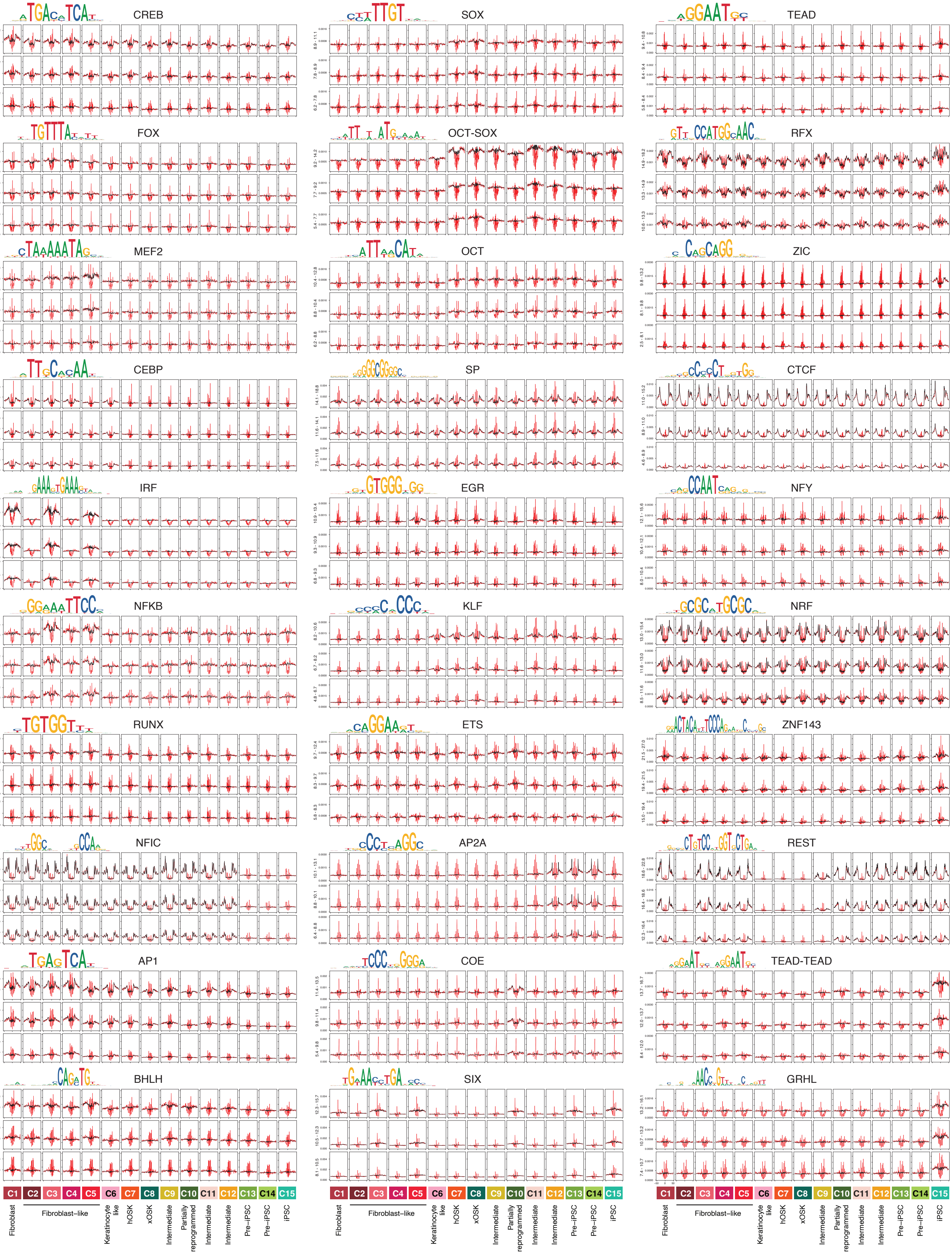
CREB
SOX
TEAD
FOX
OCT-SOX
RFX
MEF2
OCT
ZIC
CEBP
SP
CTCF
IRF
EGR
NFY
NFKB
KLF
NRF
RUNX
ETS
ZNF143
NFIC
AP2A
REST
AP1
COE
TEAD-TEAD
BHLH
SIX
GRHL
C1
C2
C3
C4
C5
C6
C7
C8
C9
C10
C11
C12
C13
C14
C15
Fibroblast
Fibroblast−like
Keratinocyte like
hOSK
xOSK
Intermediate
Partially reprogrammed
Intermediate
Intermediate
Pre−iPSC
Pre−iPSC
iPSC

**Extended Fig 3:**

Virtual footprinting for all 30 motifs obtained by inserting their instances stratified by log-odds scores into random background sequences and averaging predicted profile probability distributions with (red) and without (black) bias for each cell state.

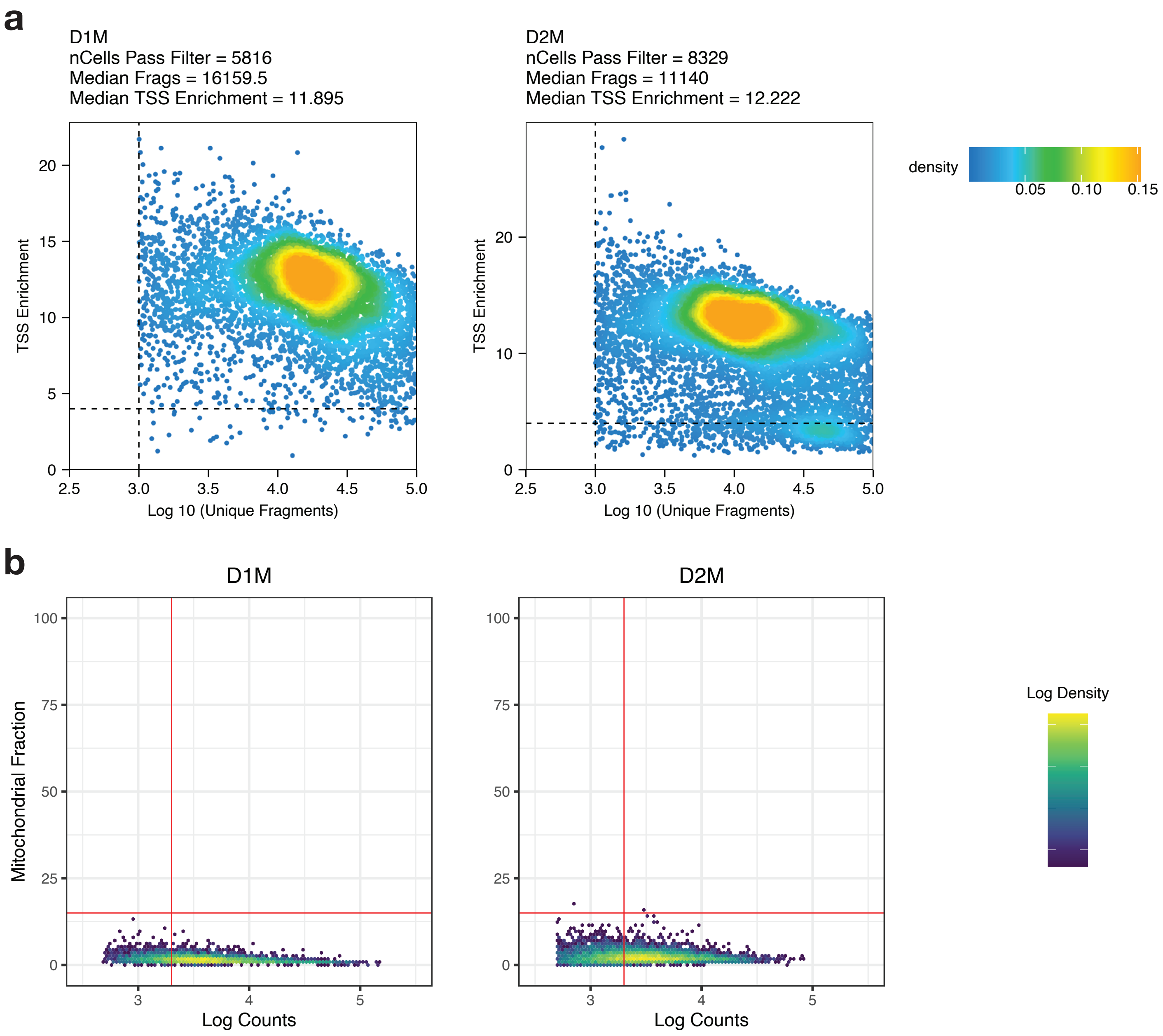


**Extended Fig 4:**
**a)** Quality control for ATAC-seq component of single nucleus multiome samples **b)** Quality control for RNA-seq component of single nucleus multiome samples
