## Supplementary Note for "Deep learning the cis-regulatory code of chromatin dynamics during cellular reprogramming"

### Base-resolution models of single-cell chromatin maps across fibroblast reprogramming reveal the interplay between transcription factor stoichiometry and motif syntax in regulating cell fate and somatic silencing

Surag Nair[1,+], Mohamed Ameen[2,3,4,5,+], Laksshman Sundaram[1], Anusri Pampari[1], Jacob Schreiber[6], Akshay Balsubramani[6], Yu Xin Wang[7,x], David Burns[7], Helen M Blau[7,8], Ioannis Karakikes[3,9,c], Kevin C Wang[4,5,10,c], Anshul Kundaje[1,6,c,#]

[+] equal contribution
[1] Department of Computer Science, Stanford University, Stanford, CA, USA
[2] Department of Cancer Biology, Stanford University, Stanford, CA, USA
[3] Cardiovascular Institute, Stanford University, Stanford, CA, USA
[4] Department of Dermatology, Stanford University, Stanford, CA, USA
[5] Program in Epithelial Biology, Stanford University, Stanford, CA, USA
[6] Department of Genetics, Stanford University, Stanford, CA, USA
[7] Baxter Laboratory for Stem Cell Biology, Stanford University, Stanford, CA, USA
[8] Department of Microbiology and Immunology, Stanford University, Stanford, CA, USA
[9] Department of Cardiothoracic Surgery, Stanford University, Stanford, CA, USA
[10] Veterans Affairs Palo Alto Healthcare System, Palo Alto, CA, USA

[x] Present address: Center for Genetic Disorders and Aging, Sanford Burnham Prebys Medical Discovery Institute, La Jolla, USA

[c] Corresponding author

#### Supplementary Note

##### Validation of reprogramming trajectory

We estimated the similarity of each cell's chromatin and expression landscape to that of canonical fibroblast cells and embryonic stem cells respectively. As expected, we observed an early loss of somatic identity and a gradual gain of pluripotency (**Fig S1a**), in line with previous observations in mouse reprogramming systems (Li et al. 2017). The chromatin-derived temporal dynamics of somatic to pluripotent identity transitions were broadly consistent with those obtained from analogous transcriptome derived similarity scores (**Fig S1b**). Reduction in expression of fibroblast markers like *COL1A1* and *FN1* was accompanied by reduced chromatin accessibility in and around their gene bodies, while changes in chromatin landscape often preceded expression of reprogramming-linked genes such as *NANOG* and *CDH1* (**Fig S1c**).

##### Peak-gene linking

To identify putative regulatory peak-gene associations, we calculated the correlation of scATAC-seq coverage and scRNA-seq expression between all pairs of scATAC-seq peaks and nearby genes across all aligned cells. 156,016 statistically significant (FDR < 0.001, t-test) peak-gene associations passed an absolute correlation threshold of 0.45, with a median 7 peaks per gene (max 97, min 0) for the top 2000 most variable genes, and 1 gene per peak (max 7, **Table S1**).

##### Fibroblast-like and keratinocyte-like trajectories

Starting from a homogeneous population of dermal fibroblast cells at day 0 (C1), heterogeneity arises immediately after OSKM induction as many cells appear to fail to initiate reprogramming

(T1, T2). Some cells remain in fibroblast-like states throughout the time-course. This fibroblast-like trajectory (T1) activates an interferon antiviral response characterized by expression of oligoadenylate synthetases (*OAS*) genes (**Fig X2a,b,c** (Gene Set 4), **X2d**) (Melchjorsen et al. 2009; Sadler and Williams 2008), and culminates in cells (C2-C5) that express higher levels of fibroblast markers such as *COL1A1* and *FN1* than the starting fibroblast population (**Fig 1g**).

Other fibroblast cells transition (T2) into a terminal population of keratinocyte-like cells (C6) that express keratinocyte markers such as *KRT14*, *KRT16* and *FLG* (Gazel et al. 2003). The population of keratinocyte-like cells (C6) along trajectory T2 expressed high levels of KM but not OS, reminiscent of a *KLF4* driven keratinocyte-like off-target cell state found in scRNA-seq analysis of mouse reprogramming (Guo et al. 2019). *KLF4* has been implicated in epidermal development (Segre, Bauer, and Fuchs 1999), and expression of *KLF4* alone in mouse embryonic fibroblasts (MEFs) can induce a keratinocyte-like fate (Guo et al. 2019).

#### Cell-state dependent OSKM concentrations

Cells that seemed to initiate reprogramming at day 2 express higher levels of all 4 factors, with *OCT4* expression levels exceeding those of the other factors (**Fig S2a,b**). Post OSKM induction, day 2 cells split into either hOSK (C7, median expression across cells for OSKM 1863, 583, 893, 1023 TPM respectively) or xOSK (C8, OSKM expression 7217, 2194, 2440, 1196 TPM respectively) states (**Fig S2a**). Median expression levels for day 2 cells in the xOSK state were 4-fold higher for *OCT4* and 8-fold higher for *SOX2* compared to iPSC levels (*OCT4*: 1740 TPM, *SOX2*: 271 TPM, negligible KM ~ 0 TPM) (**Fig S2a**).

While premature withdrawal of OSKM expression can stall the progress of cells in a partially reprogrammed state (Tanabe et al. 2013; Polo et al. 2012), eventual silencing of transgenes is required for maturation of Pre-iPSC cells (Golipour et al. 2012). Compared to day 2 cells in xOSK state, day 10 cells in the Intermediate state C12 along the reprogramming trajectory T3 were down nearly 10-fold for OSK (OSKM expression 833, 175, 176, 750 TPM respectively, **Fig S2a**) and the final population of iPSCs was transgene-free. Transgene expression levels were higher along states on the primary reprogramming trajectory (T3) compared to the partially reprogrammed trajectory (T4) (OSKM expression 305, 77, 64, 522 TPM respectively for day 10 cells in Intermediate state C9).

#### TF2G Networks for FN1 and JUN

TF2G networks of the locus around the fibroblast-specific gene *FN1* implicates somatic TFs AP-1, RUNX and TEAD as its primary regulators in fibroblasts. On induction of OSKM, the *FN1* locus rapidly loses accessibility at multiple enhancers and by Day 4 its expression is reduced more than 10 fold (**Fig S5a,b**). Inferred TF2G networks regulating *JUN*, a member of the AP-1 subunit, implicate motifs of TEAD and KLF/SP factors as well as autoregulatory feedback involving gradual loss of AP-1 dependent enhancers over the course of reprogramming (**Fig S5c**) (Angel et al. 1988).

#### Keratinocyte-specific and Stable peak sets

A set of keratinocyte-specific peaks (K1, 28388 peaks) is highly accessible in the keratinocyte-like population (C6), linked with 781 genes including keratinocyte markers such as *KRT16* and *KRT14*, and enriched for the GO term cornification (**Fig 4a-d**). As expected,

Keratinocyte-like specific peaks (K1) are enriched primarily for KLF motifs but not OCT-SOX. Finally, constitutively accessible peaks (Stable S1-3, 80873 peaks) are localized near both fibroblast and iPSC genes, as well as housekeeping genes such as *GAPDH* and *RPS11*.

#### Late transient (COC/L) peaks and role of *TFAP2C*

Late Transient peaks (COC/L1-5) attained maximum accessibility between days 4-14, and then close in iPSCs, likely arising due to secondary effects of OSKM overexpression. COC/L peaks are relatively depleted for O binding but displayed high scores for TFAP2 motifs (**Fig 4d, e (iii)**), corresponding with a hybrid naive-primed pre-iPSC state (**Fig S9b, Supplementary Note**).

*TFAP2C*, which is a pioneer factor, is the only *TFAP2* family TF expressed during reprogramming (**Fig X2d**). COC/L peaks are linked to pre-implantation genes such as *DPPA2/3/5* and *DNMT3L* that are preferentially expressed in Pre-iPSC cells (**Fig X2d**). This is consistent with the observation that elevated levels of *KLF4* in Pre-iPSCs relative to iPSCs activate *TFAP2C* to drive cells to a transient naive, pre-implantation-like state (Liu et al. 2020; Pastor et al. 2018; Tan et al. 2011; Cacchiarelli et al. 2015; Gafni et al. 2013; Takashima et al. 2015). The chromatin accessibility landscape of Pre-iPSC cells resembles a hybrid naive-primed state (**Fig S9b**). The hybrid naive-primed state likely facilitates media-dependent exit into either naive or primed iPSC state.

#### Support for an ‘enhancer selection’ model

Different combinations of motifs are engaged at different time points across the time course via cooperative binding of OSK with other TFs that are expressed at specific time points. For example, at the CO/E1 peak chr13:47816815-47817315, the early gain of accessibility in the xOSK state (C8) is attributed to OCT-SOX and KLF motifs. Over the course of reprogramming, contribution of the KLF motif diminishes while a predictive ZIC motif instance emerges in the iPSC state, concomitant with late expression of multiple *ZIC* TFs (**Fig 4e (i), Fig X2d**), in line with the late stage 'enhancer selection' model proposed in Chronis *et al.* (Chronis et al. 2017).
