## Supplementary material for "Deep learning the cis-regulatory code of chromatin dynamics during cellular reprogramming": Methods

### Transcription factor stoichiometry, motif affinity and syntax regulate single-cell chromatin dynamics during fibroblast reprogramming to pluripotency

Surag Nair[1,+], Mohamed Ameen[2,3,4,5,+], Laksshman Sundaram[1], Anusri Pampari[1], Jacob Schreiber[6], Akshay Balsubramani[6], Yu Xin Wang[7,x], David Burns[7], Helen M Blau[7,8], Ioannis Karakikes[3,9,c], Kevin C Wang[4,5,10,c], Anshul Kundaje[1,6,c,#]

[+] equal contribution
[1] Department of Computer Science, Stanford University, Stanford, CA, USA
[2] Department of Cancer Biology, Stanford University, Stanford, CA, USA
[3] Cardiovascular Institute, Stanford University, Stanford, CA, USA
[4] Department of Dermatology, Stanford University, Stanford, CA, USA
[5] Program in Epithelial Biology, Stanford University, Stanford, CA, USA
[6] Department of Genetics, Stanford University, Stanford, CA, USA
[7] Baxter Laboratory for Stem Cell Biology, Stanford University, Stanford, CA, USA
[8] Department of Microbiology and Immunology, Stanford University, Stanford, CA, USA
[9] Department of Cardiothoracic Surgery, Stanford University, Stanford, CA, USA
[10] Veterans Affairs Palo Alto Healthcare System, Palo Alto, CA, USA

[x] Present address: Center for Genetic Disorders and Aging, Sanford Burnham Prebys Medical Discovery Institute, La Jolla, USA

[c] Corresponding author

#### Methods

### 1. Experimental methods

#### 1.1. Fibroblast isolation and culture

Skin punch biopsy (2 × 2 mm) was performed on patients following Stanford Institutional Review Board (IRB) approval and Stem Cell Research Oversight (SCRO) approval at Stanford University and under informed consent. The skin biopsy was manually minced with scissors and incubated in collagenase type IV (Life Technologies) for four hours. Large debris was removed and fibroblasts were centrifuged and resuspended in DMEM with Glutamax (Invitrogen) supplemented with 10% FBS. Fibroblasts were expanded in culture at 37 °C, 20% O2, and 5% CO2 in a humidified incubator over two weeks and were frozen at passages 3 and 4 until required for reprogramming.

#### 1.2. Reprogramming

Sendai virus reprogramming was performed according to the Cytotune Sendai Viral reprogramming protocol (Invitrogen). Briefly, fibroblasts were grown in DMEM with 10% fetal bovine serum and were trypsinized the day before infection and 40,000 fibroblasts were plated into a Matrigel-coated well of a 24-well plate. The following day, Sendai virons containing *POU5F1*, *SOX2*, *KLF4* and *cMYC* genes were added to the well at MOI=5-5-3 (KOS MOI=5, hc-Myc MOI=5, and hKlf4 MOI=3), and the tissue culture plate was spun for 15 min at 800g. For 7 days after viral infection, fibroblasts were grown in fibroblast media and the media was refreshed every other day.  On day 7 after infection, fibroblasts were cultured in E8 media for 12 additional days.

#### 1.3. Single cell RNA-seq data generation

We profiled reprogramming of human fibroblasts at eight time points separated by 48 hours. Trypsinization was carried out for 5 min for all time points, followed by addition of fibroblast medium to inactivate the trypsin. Following centrifugation, cells were washed with 1X PBS containing 0.1% bovine serum albumin. The cells were then passed through a 40 micron filter to remove cell debris. Using a cell counter, the number of cells and viability were calculated to a final concentration of ~1000 cells/ul. With a target of 10000 cells per GEM reaction, single cell RNA-seq libraries were generated at each time point using 10X Genomics Single Cell 3′ Reagent Kits v3 according to manufacturer's instructions. We sequenced the pooled libraries using the Illumina NovaSeq 6000.

#### 1.4. Single cell ATAC-seq data generation

The cells were dissociated using Tryple Express and resuspended in fibroblast medium. The Cells with viability >90% were washed in ice-cold ATAC-seq resuspension buffer (RSB, 10 mM Tris pH 7.4, 10 mM NaCl, 3 mM MgCl2), spun down, and resuspended in 100 mL ATAC-seq lysis buffer (RSB plus 0.1% NP-40 and 0.1% Tween 20). Lysis was allowed to proceed on ice for 5 min, then 900 mL RSB was added before spinning down again and resuspending in 50 mL 1X Nuclei Resuspension Buffer (10x Genomics). A sample of the nuclei was stained with Trypan Blue and inspected to confirm complete lysis. Nuclei were processed using a 10X chromium single-cell ATAC-seq kit (version V1, 10X Genomics) at the Stanford Functional Genomics Facility (SFGF). All samples were sequenced using the NovaSeq 6000.

##### 1.5. Multiome data generation

For multiome single cell data, human fibroblast reprogramming was repeated at MOI=5-5-3 (KOS MOI=5, hc-Myc MOI=5, and hKlf4 MOI=3). Day 1 and 2 post Sendai transduction, the nuclei were prepared as above for ATAC-seq with minor changes. Specifically, 0.01% digitonin was added to the lysis buffer, and 2 U/µL RNase inhibitor (Roche) was added to all nuclei preparation buffers. After nuclei preparation, droplets and single cell libraries were prepared using the Single Cell Multiome ATAC + Gene Expression kit (10x Genomics).

#### 2. Computational methods

##### 2.1. Single-cell ATAC-seq processing

###### 2.1.1. Mapping

We used Chromap (commit 6e97125b9 https://github.com/haowenz/chromap/tree/6e97125b9) (Zhang *et al.*, 2021) to perform barcode correction, alignment and filtering for each of our samples. We built an index using the default Chromap parameters (kmer length=17, window size=7) for the hg38 build. We aligned each of our samples with the prepared index and the flags "--preset atac --drop-repetitive-reads 4 -q 0". This allows multi-mappers that align at up to 4 positions and assigns them to one position, which avoids gaps in alignment at regulatory elements that can not be aligned to with unique reads. We used the 10x ATAC-seq barcode whitelist (737-cratac-v1.txt.gz, copy available at https://github.com/caleblareau/errorcorrect_10xatac_barcodes/blob/5b9380b92/737K-cratac-v1.txt.gz) after reverse-complementing each barcode. The output of this step is fragment files, one for each sample. Each fragment file contains five columns, the first 3 correspond to chromosome, start and end of the fragment, the fourth contains the corrected barcode and the final contains the number of read pairs associated with that fragment.

##### 2.1.2. Quality control

We used ArchR v1.0.1 (Granja *et al.*, 2021) for initial quality control. We used the “createArrowFiles” function which takes as input all the fragment files from the mapping stage. In addition, we run the “addDoubletScores” function with parameters “k=10, knnMethod=’UMAP’, LSIMethod=1”. These methods return quality control measures for each barcode as well as doublet enrichment scores. We use thresholds: total fragments >= 1000, TSS enrichment >= 6 and doublet enrichment <= 1.25 to filter barcodes. This step returns a total of 65,357 viable barcodes across all samples.

##### 2.1.3. Dimensionality reduction, clustering, peak calling and UMAP

We used SnapATAC (Fang *et al.*, 2021) for dimensionality reduction, clustering and constructing the UMAP. We added bug fixes to the original snapATAC codebase. The code can be accessed at https://github.com/kundajelab/SnapATAC/tree/4cb43f7. We used a two-stage procedure that is in part based on tutorials provided with the SnapATAC repository. In the first stage, we perform dimensionality reduction and clustering based on counts in 5kb tiled genomic bins. After refining the clusters, we call peaks for each of the individual clusters treating them as pseudo-bulk ATAC-seq, and aggregate them to obtain a set of peaks. In the second stage, the dimensionality reduction and UMAP are recomputed based on the counts in peaks. The specifics are described in more detail below.

We first analyzed a binary counts matrix over tiled 5kb genomic bins. Bins overlapping with the hg38 blacklist (https://www.encodeproject.org/annotations/ENCSR636HFF/) (Amemiya *et al.*,

2019) were omitted. We removed the top 5% of bins with the most cells. This gave a total of 543,838 bins. We sampled 1200 cells from each of the 9 timepoints as landmark cells for diffusion maps (runDiffusionMaps method with num.eigs=50). The remaining cells were projected using the runDiffusionMapsExtension method. We called the runKNN method with eigs.dim=1:10, k=15 in order to construct the kNN graph from the first 10 eigenvectors. The runViz method with method="umap", eigs.dim=1:10 was used to create an initial UMAP. We performed clustering with the runCluster function with arguments louvain.lib="leiden", resolution=2.25. The clusters were manually adjusted to obtain a total of 16 clusters. One of the clusters contained a clump of cells from across all samples and had low overall TSS enrichment, which we omitted from further analysis. We also removed a small population of cells colocated with the clump and detached from its own cluster in UMAP space, which are likely doublets. 82 cells in the iPSC-specific cluster that were not from the iPSC sample were also omitted. These cells, primarily from D14, were not colocated with the iPSC cluster in the UMAP but instead with pre-iPSC cells. These omissions reduce the number of cells from 65,357 to 62,715.

We next extracted fragments from each cluster and ran the bulk ENCODE ATAC-seq pipeline v1.9.1 (Hitz *et al.*, 2023). The pipeline returns peak calls and signal tracks for each of the clusters. The displayed cell state specific signal tracks throughout the paper correspond to -log *p*-value tracks derived from the pipeline outputs, with y-axes fixed between 2 and 40. In order to obtain a combined peak set, we used an algorithm similar to ArchR's approach for combining peaks. We extended each peak from its summit by 250bp on both sides. We considered peaks across clusters in order of their increasing q-values and accepted peaks that did not overlap with any previously accepted peak. This resulted in a peak set with a total of 530,910 500bp peaks.

For the second stage, we constructed a peak by cell matrix with the combined peak set. The dimensionality reduction followed the same procedure as in the first stage, with identical steps and parameters, with the exception of an additional filtering step where cells with fewer than 500 peaks (after binarizing) were discarded resulting in 62,599 cells. The resulting UMAP was highly similar in structure to the UMAP constructed from the 5kb bin matrix in stage 1. We retained cluster assignments from stage 1.

###### 2.1.4. ATAC-seq derived gene scores

We used the GeneScoreMatrix generated by ArchR for ATAC-based gene scores. For visualization, we smoothed the matrix using the nearest neighbor matrix (k=15).

###### 2.1.5. Peak set clustering

We clustered peaks into peak sets to capture shared dynamics across cell states. We initially split all peaks into 4 possible states based on overlap with fibroblast and iPSC peaks (Fibroblast only, iPSC only, shared or neither). Peaks were clustered separately within each of the 4 states. In order to perform the peak clustering, we first clustered cells with Leiden clustering with high resolution (resolution=5). By doing so, we can capture fine scale dynamics while also mitigating the sparsity of scATAC-seq at single-cell resolution. This split the cells into 71 clusters with a median of 872 cells in each (min 150, max 1823). We aggregated counts for each cluster pseudo-bulk and constructed a reduced peak by cluster pseudo-bulk counts matrix. The counts were normalized with the estimateSizeFactorsForMatrix function from DESeq2 (Love *et al.*, 2014). We then log2 transformed the matrix with a pseudocount of 1. We concatenated the z-normalized matrix (normalized with the scale function) with the log normalized counts matrix. The main motivation behind using both log normalized counts and scaled values is that both the absolute and relative accessibility values are important-- we would like to discriminate between two peaks with the same accessibility dynamics but different absolute chromatin accessibility. E.g., two peaks may increase in accessibility over reprogramming, but one may start from closed chromatin and become accessible, while the other may start from accessible chromatin and increase accessibility further.

We perform k-means clustering for each of the 4 states separately. Clustering is performed using the ClusterR package (Mouselimis *et al.*). We used the MiniBatchKmeans function with batch_size=1000, num_init=50 and variable number of clusters for each of the 4 states- 3 for fibroblast only, 4 for iPSC only, 3 for shared and 14 for neither (transient). We then manually adjusted the 14 peak sets obtained for transient peaks, discarded a peak set with low overall accessibility and low variability, and ended with 20 peak sets in all.

#### 2.2. Single-cell RNA-seq processing

##### 2.2.1. Mapping and quantification

We used 10x Genomics Cell Ranger v6.0.2 (Zheng *et al.*, 2017) for read mapping and quantification to obtain the counts matrix. We used the GRCh38 2020-A reference. For each sample, we ran Cell Ranger count with expect-cells=10000. After processing each sample separately, we aggregated across samples using the Cell Ranger aggr command with --normalize=none. This ensures no reads are dropped by subsampling when aggregating counts across samples. The filtered_feature_bc_matrix is used for all downstream analyses.

##### 2.2.2. Quality control

We primarily used Seurat v3.1.5 (Stuart *et al.*, 2019) for scRNA-seq analyses. The initial filtered_feature_bc_matrix matrix contained 86,089 barcodes across all samples. We applied the following filters for quality control for each barcode: >200 unique detected genes, total UMIs >2000 and <100,000 and mitochondrial fraction <15%. In addition, we observed that some cells, primarily from Day 2, exhibited a high total fraction of OSKM UMIs. More than 10% of Day 2 cells had >10% OSKM fraction, and 1% had >50% OSKM. We applied an additional filter to remove cells with >50% OSKM fraction.

We used DoubletFinder v2.0.3 (McGinnis *et al.*, 2019) to detect and remove doublets. After applying the filters described above, we ran DoubletFinder for each sample individually. We processed each sample and performed normalization (NormalizeData), scaling (ScaleData), PCA (runPCA) on 2000 variable genes (FindVariableFeatures) and UMAP (runUMAP) with the first 10 principal components. We then ran the doubletFinder_v3 function with PCs=1:10, pN=0.25, pK=0.01 and nExp = 0.15 x number of barcodes, which eliminated 15% of cells from each sample. After filtering and doublet removal, we ended with 59,378 barcodes across all time points.

##### 2.2.3. Dimensionality reduction and UMAP

We first normalized the counts data (NormalizeData) for all barcodes. We then subsampled 2630 barcodes from each time point, giving 23,670 barcodes in total. 2630 was chosen since it was the number of barcodes in Day 2 samples, which was the smallest number among the time points. In contrast, 10,443 barcodes passed quality control for the Day 10 sample. Thus, by subsampling prior to computing the principal components, we avoid being overly biased by samples with more barcodes. We performed scaling (ScaleData) and PCA (runPCA) on 2000 variable genes (FindVariableFeatures) on the subsampled matrix. We then applied the scaling and PCA transformation on the remaining cells. Finally, we called the runUMAP function with dims=1:10 in order to obtain the UMAP layout for the scRNA-seq cells.

##### 2.2.4. Unbiased Gene set clustering

We clustered genes into gene sets from scRNA-seq to capture genes with shared co-expression patterns across cell states. The overall approach follows clustering of scATAC peaks into peak sets as described above. In order to perform the gene clustering, we first clustered scRNA cells with high resolution=1. This split the cells into 35 clusters with a median of 1486 cells in each (min 135, max 5510). We aggregated counts for each cluster pseudo-bulk and constructed a reduced gene by cluster pseudo-bulk counts matrix. The counts were normalized with the estimateSizeFactorsForMatrix function from DESeq2. We then log2 transformed the matrix with a pseudocount of 1 followed by z-scaling. We perform k-means clustering. Clustering is performed using the ClusterR package. We used the MiniBatchKmeans function with batch_size=1000, num_init=50 and 15 clusters. We then manually reordered the 15 gene sets.

##### 2.2.5. Endogenous OSKM quantification

To quantify per barcode endogenous OSKM UMI counts, we relied on the fact that Sendai mRNA transcripts terminate at stop codons, whereas endogenous transcripts contain the 3' UTR. For each of OSKM, we constructed a reference distribution for Sendai-derived 5' read ends from day 2 pseudo bulk (assuming no endogenous transcripts), and a reference distribution for endogenous 5' read ends from iPSC pseudo bulk (assuming no Sendai transcripts). We modeled each barcode as a mixture of the two 5' end distributions with unknown weights and used an Expectation-Maximization based iterative algorithm to estimate the fraction of endogenous transcripts for each barcode.

###### 2.2.6. Sendai gradient "velocity"

This plot depicts arrows pointing in the direction of decreasing total Sendai expression over the scRNA-seq UMAP. For each cell, we took its 14 nearest neighbors in the kNN graph. To obtain a "gradient" on the UMAP, we computed the difference between the UMAP coordinates for the cell with each of its nearest neighbors and normalized them to be unit vectors. We then averaged these vectors, weighting them the difference in total OSKM Sendai expression between the cell-pair, to obtain a "Sendai gradient velocity" for each cell (distinct from typical RNA velocity). We discretized the UMAP and averaged the velocity in each bin. We removed regions with low total Sendai expression and plotted the resulting arrows using matplotlib's streamplot function.

##### 2.3. scATAC and scRNA integration

We assigned labels to scATAC-seq clusters and then transferred the labels to scRNA-seq cells. To achieve this, we called the FindTransferAnchors function in Seurat using the reference as the scATAC-seq gene score matrix computed using ArchR and query as the scRNA-seq expression matrix. CCA reduction was used on a set of variable genes which had gene scores available. The function was called together for cells across all samples. The scRNA-seq cluster score predictions were made using the TransferData function with weight.reduction set to PCA reduction of scRNA-seq data. This returned a cell by cluster matrix which had predicted scores for each scRNA-seq cell across all clusters. We created a masking matrix of samples (Days 0,2...14 and iPSC) by cluster in which any (cluster, sample) pair that contained fewer than 2% cells of the sample in scATAC-seq data was set to 0. We applied this mask on the predicted cluster scores for scRNA-seq cells in order to avoid assigning cells to clusters that are unlikely based on their sample, e.g. no cells from the Day 0 fibroblast sample should be assigned the iPSC cluster label. The label with the highest predicted score after masking was chosen as the cluster for each scRNA cell.

To get a common representation of both scRNA and scATAC cells, we again used the FindTransferAnchors function in Seurat this time using the reference as scRNA-seq expression matrix and query as scATAC-seq gene score matrix computed using ArchR. We then extracted the L2 normalized CCA embeddings and created a UMAP with 30 CCA dimensions, but observed that the scRNA and scATAC cells were not sufficiently mixed. Thus, we ran Harmony (Korsunsky *et al.*, 2019) to correct for assay-specific effects using the RunHarmony function on the L2 normalized CCA embedding. We used 30 components of the Harmony embeddings for downstream applications and created a UMAP with both scRNA and scATAC cells. To obtain the nearest scRNA neighbor of a scATAC cell, we find the nearest neighbor in the Harmony space, subject to the constraint that both cells must be assigned the same cluster label.

To link peaks to genes, we next used ArchR's addGeneIntegrationMatrix function followed by the addPeak2GeneLinks function. We use an absolute correlation cutoff of 0.45 and FDR cutoff 1e-4.

#### 2.4. OSKM analysis

##### 2.4.1. <u>ChromVAR</u>

We used HOCOMOCO (Kulakovskiy *et al.*, 2018) motifs SOX2_HUMAN.H11MO.0.A, KLF4_HUMAN.H11MO.0.A, MYC_HUMAN.H11MO.0.A and JASPAR (Castro-Mondragon *et al.*,

2022) motifs POU5F1_MA1115.1 and OCT-SOX_MA0142.1. We ran ChromVAR (Schep *et al.*, 2017) with default parameters on the binarized peaks x cells matrix

###### 2.4.2. OSKM expression states

We clustered cells based on log expression values of OSKM. We used the MiniBatchKmeans function of ClusterR library with clusters=8, batch_size=100, num_init=50, which bucketed each cell into one of 8 states based on the pattern of OSKM expression. We then manually labeled these states based on the average expression of OSKM in each state.

##### 2.5. Trajectory analysis

###### 2.5.1. PAGA analysis

We used PAGA v1.4.4.post1 (Wolf *et al.*, 2019, 2018) to construct a cell state by cell state connectivity graph on the scATAC data. We first ran pp.neighbors with n_neighbors=15 and used the first 10 components of the diffusion map, followed by the tl.paga function with default arguments. We computed pseudotime values for cells by running the tl.diffmap function followed by the tl.dpt function. We used two different root cells to compute two pseudotime values per cell. The first root was a fibroblast Day 0 cell, and the second was a cell from the xOSK Day 2 state. By treating fibroblast cells as root, we can use the derived pseudotime values for the cells that retain fibroblast-like characteristics throughout the time course. On the other hand, using the xOSK cells as root, we can use the derived pseudotime values for reprogramming cells. The underlying assumption is that the cells that express ectopic OSKM at high levels are transported between Day 0 and Day 2 to a distinct xOSK state for which we do not observe cells along the transition due to insufficient temporal resolution of samples in these early stages. As a result, xOSK Day 2 cells can be treated as root cells for reprogramming cells from subsequent time points.

##### 2.5.2. Trajectory construction

We draw three main trajectories on the UMAP, which correspond to the reprogramming trajectory (clusters 8, 11-14), partially reprogramming trajectory (clusters 7, 9, 10) and fibroblast-like trajectory (clusters 1-5). For the first two, we use pseudotime values using xOSK cell as root, and pseudotime values derived using fibroblast cell as root for the fibroblast-like trajectory. For each of the trajectories, we drew lines on the UMAP. We ordered all cells belonging to a trajectory by pseudotime, and used the R function smooth.spline on their x and y coordinates separately with df=200 and spar=1 for smoothing, followed by ggplot's geom_path. In addition, we manually added 3 dashed lines to the UMAP, representing unobserved transitions in our data. The 3 unobserved transitions are the initial transitions from Day 0 fibroblasts to Day 2 xOSK and keratinocyte-like cells, and a transition from Day 14 pre-iPSC cells to iPSC cells.

#### 2.6. Gene set enrichment analyses

We used the R package gprofiler2 (Raudvere *et al.*, 2019) for gene set enrichment analyses which is an interface to the g:Profiler portal (version e104_eg51_p15_3922dba). For every peak set, we considered genes that were linked to at least 2 peaks in that peak set and ranked them in descending order of the number of peaks they were linked to in that peak set. We used the gost function with organism="hsapiens" and ordered_query=T.

#### 2.7. ChromBPNet: a deep learning sequence model of base-resolution chromatin accessibility profiles

##### 2.7.1. Code

The code for model training, evaluation and interpretation used in this publication is available at https://github.com/kundajelab/scATAC-reprog/tree/master/src/chrombpnet. However, further development of this repository has been deprecated. Instead, users who wish to use ChromBPNet for training and interpreting models on custom datasets should instead use the latest up-to-date version of ChromBPNet at https://github.com/kundajelab/chrombpnet.

##### 2.7.2. Base-resolution coverage tracks of cell state resolved pseudobulk scATAC-seq profiles

ChromBPNet models of each cell state are trained to map local DNA sequence windows to their associated base-resolution coverage profiles of pseudobulk scATAC-seq data aggregated across all cells belonging to the cell state. For each cell state, we first extracted fragments from the scATAC-seq fragment files associated with cell barcodes that are assigned to the cell state. We filtered out reads mapping to chrM and adjusted the 5' ends of reads to get an effective shift of +4 on "+" strand and -4 on "-" strand to account for Tn5 transposase offset. Most ATAC-seq pipelines typically recommend a +4/-5 shift. However, a +4/-4 shift in fact aligns the Tn5 offset on the two strands exactly which is critical for accurate modeling of Tn5 sequence preferences for coverage tracks aggregated across both strands. We then proceeded to convert the filtered and shifted 5'-ends (cut-sites) from both strands into base-resolution coverage tracks (BigWig format). To do so, we converted the adjusted fragment file to tagAlign files, where both reads of each fragment (fragment) are used in a strand specific manner. The tagAlign file was then converted to a bedGraph coverage file with the "bedtools genomecov -5 -bg -g” command after sorting the input files which counts 5’ ends of reads from both strands at each base. The obtained bedGraph was sorted and converted to a BigWig file with the “bedGraphToBigWig” command.

##### 2.7.3. Enriched signal regions and background regions selection for training ChromBPNet in a chromosome hold-out cross-validation set up

For each cell state, we obtain the reproducible sets of pseudo-bulk scATAC-seq peaks based on overlap of MACS2 peaks (*p*-value < 0.01) called on pseudo-replicates, which is an output of the ENCODE ATAC-seq pipeline (v1.9.1) https://github.com/ENCODE-DCC/atac-seq-pipeline. We merged any overlapping peaks (“bedtools merge”) and created 3000 bp windows from the center of each merged peak. 3000 bp was chosen to account for the 2000 bp input of the model and to allow for 500 bp of jittering during training on either side. Note that there may be some overlap between the regions. We also removed any regions overlapping the GRCh38 Exclusion List (blacklist) of problematic regions (Amemiya *et al.*, 2019) https://www.encodeproject.org/annotations/ENCSR636HFF/. This formed the set of enriched signal regions for each cell state.

We split the hg38 genome into 3000 bp regions. For each cell state, we created a set of background regions, equal in number to the enriched signal regions, that did not overlap with either any enriched signal region, any reproducible peaks or any blacklisted region. In addition, the background regions were chosen to match the GC content distribution of the enriched signal region set as closely as possible. The background regions are non overlapping. We used a 10-fold chromosome hold-out cross-validation setup. The train, validation and test chromosomes corresponding to each of the 10 folds can be found at the following Zenodo archive https://doi.org/10.5281/zenodo.8299710.

##### 2.7.4. ChromBPNet overview

A major consideration when modeling base-resolution ATAC-seq coverage profiles as a function of sequence is that the Tn5 transposase in the ATAC-seq assay has a strong sequence preference for cleavage sites. There can be an order of magnitude difference between sequence preference for even two adjacent bases. Therefore, local cutting patterns are strongly determined by the underlying sequence independent of regulatory sequence drivers of chromatin accessibility. Accounting for this sequence preference, which we refer to as Tn5 bias (or simply bias), is essential to disentangle regulatory sequence features, decipher bias-corrected ATAC-seq footprints of TFs and improve overall model interpretability. To learn and correct for Tn5 sequence bias, ChromBPNet fits a sequence model to base-resolution ATAC-seq coverage profiles in two stages.

In the first stage, a neural network (the “bias model”) is trained to explicitly learn base-resolution Tn5 insertion coverage in background regions (not overlapping any peak regions) as a function of sequence. This model maps 2000 bp DNA sequence to a 2000 length vector of logits (log probabilities) of Tn5 insertions per position, as well as a scalar count output that corresponds to bias dependent counts in background regions expected to be devoid of transcription factor activity. In the second stage, we train a cell state specific model (the “accessibility model”) which predicts the observed base-resolution ATAC-seq signal profiles within peaks. This model takes as input 2000 bp DNA sequence, as well as the predictions of the bias model (Tn5 bias logits and scalar count) on the same DNA sequence. The accessibility model initially outputs (latent) de-biased profile logits and scalar count. These de-biased logits and count are additively combined with the bias model’s logits and predicted counts to get the final model output profile logits and predicted count. The accessibility model is trained on the observed ATAC-seq signal profiles within peak regions and background regions. After training, the bias model is “unplugged” which effectively regresses out the bias from the predictions. The de-biased profile logits and counts are analyzed and interpreted to reveal cell state specific insights.

##### 2.7.5. Bias model architecture

The bias model predicts Tn5 sequence preference over a given input sequence. It takes as input a 2000 length sequence input, and outputs profile logits (log probabilities) over the 2000 bases as well as log counts over the input sequence. We use the BPNet model architecture in this work, which is nearly identical to the architecture in the BPNet paper (Avsec *et al.*, 2021). Specifically, the bias model takes as input a 2000 length input sequence as a one-hot encoded matrix. This is passed through an initial convolution layer with 256 filters and kernel size 25. The output is then passed through 9 successive dilated convolution layers, each with 256 filters, kernel size 3 and dilation rate 2^i (i for layers from 1 to 9). All convolutions have padding=“same” and activation=”relu”. For the 9 successive dilated convolution layers, the input to the layer is added to its output akin to a residual layer. The output of the final dilated convolution layer (termed bottleneck output) is passed through an additional convolution layer with kernel size 25, padding=”same”, and one output channel, which is flattened to obtain the profile logits output of the model. Separately, the bottleneck output passed through a global average pooling layer, followed by a fully connected layer with 1 output, to obtain the log counts output of the model.

##### 2.7.6. Accessibility model architecture

The accessibility model (ChromBPNet model) predicts the ATAC-seq profile for a given cell state over a given input sequence. In addition to 2000 length sequence input, it also takes as input the bias model predicted logits and log counts over the same sequence. The accessibility model processes the sequence input with an architecture identical to the bias model's. This returns (unobserved) de-biased profile logits and log counts. The final profile prediction of the accessibility model is simply the sum of the de-biased profile logits and the bias model predicted logits. The final log counts prediction of the accessibility model is obtained by combining the de-biased log counts output with bias model predicted log counts through a log-sum-exp operation, i.e. the accessibility model's predicted log counts are the logarithm of the sum of counts predicted by the bias model and de-biased counts.

###### 2.7.7. Bias model training

The bias model is trained to map a 2000 bp input sequence to per position log probabilities predictive of Tn5 insertion preference and log of total counts that can be attributed to background Tn5 activity. In order to learn the implicit Tn5 sequence preference, we utilize non-peak background regions. While a large fraction of reads fall within peak regions, a significant number of reads fall outside peak regions. We assume that the reads mapping outside peaks correspond to background Tn5 activity devoid of influence of transcription factors. This is similar to the approach adopted by TOBIAS (Bentsen *et al.*, 2020). In addition, to ensure no TF-dependent signal is picked up when training the bias model, the background regions are further thresholded by the total number of Tn5 insertions in the region. We use a conservative threshold of half the minimum number of total counts observed in any peak (positive) region.

We use a multinomial negative log likelihood (MNLL) for the profile output and a mean square error (MSE) loss for the log counts output of the model. The relative loss weight used for the counts loss is 0.1 x mean per region total counts. During every epoch, the training examples were randomly jittered by up to 100bp on either side, and a random half of the examples were reverse-complemented (the corresponding profile label is also reversed). Adam optimizer is used with default parameters, with an early stopping criteria of 5 consecutive epochs without validation loss improvement. All analyses including model training and interpretation were performed using Tensorflow v1.14.0 and Keras v2.2.4.

We observed that a Tn5 bias model trained on one cell state worked reliably for Tn5 bias correction in other cell states as well. Thus, we utilize a single Tn5 bias model that was trained on the fibroblast cell state (C1). Furthermore, we performed standard interpretation and motif discovery (described in the Interpretation and TF-MoDISco sections) on the trained Tn5 bias model and ensured that the model did not discover any TF motifs. We adjust the log counts output of the bias model for each new cell state by adding a constant (in log space, which is equivalent to scaling the counts) to account for variable sequencing depth, such that it minimizes the squared error on all background regions of the new sample. This is called the cell state specific adjusted bias model.

##### 2.7.8. Accessibility model training

The accessibility model is trained on peaks and background regions. The loss and optimization routine is equivalent to that of the bias model. During every epoch, the training examples are jittered by up to 500 bp on either side, and a random half of the sequences are reverse-complemented. The adjusted bias model is called to predict the bias logits and logcounts for the jittered and/or reverse-complemented sequences, which are fed as input to the accessibility model. By explicitly providing the Tn5 bias logits and log counts as an input which is added into the accessibility model's output, the accessibility model learns the residual accessibility signal that is independent of the transposase's sequence preference. During inference, we can make profile predictions without providing the Tn5 bias logits and log counts, and hence obtain bias corrected profile predictions.

##### 2.7.9. Model Evaluation

We evaluated the counts and profile outputs of each model on the test set for each cross-validation fold. For counts, we computed the Pearson correlation between the predicted log counts and the observed log counts over only peaks, and separately over peaks and background regions. For the profile predictions, we first converted the predicted logits for every sequence to a predicted probability distribution through a softmax operation. For each peak sequence, we then computed Jensen-Shannon Divergence (JSD) between the predicted probability distribution and the observed read distribution. We then computed the median of these values across all peak regions as a metric of profile performance. Note that lower is better for the median JSD. We estimated a lower bound (i.e. best-case bound) by computing the JSD between pseudoreplicates of the observed signal (each with half the read depth of the original signal) in an identical fashion. We also obtained an upper bound (i.e. worst-case bound) by computing the median JSD of the bias model predicted profile distribution against the observed read distribution. Note that it is difficult to compare profile performance across cell states as the JSD depends on read depth of the samples, which determines the ground truth.

##### 2.7.10. Estimating sequence contribution scores and *de novo* motif discovery

We use the models to estimate the contribution of each base pair in any sequence of interest to the predicted outputs as outlined previously in (Avsec *et al.*, 2021). For each ChromBPNet model, we use the DeepSHAP implementation of the DeepLIFT algorithm (https://github.com/kundajelab/shap/tree/29d2ffab) (Shrikumar *et al.*, 2017; Lundberg and Lee,

2017) to derive base-resolution contribution scores across any input sequence of interest. DeepSHAP is run using 20 dinucleotide shuffled versions of the input sequence as references, and computed separately with respect to the count (called counts contribution scores) and profile (called profile contribution scores) outputs of the model. The profile contribution scores are more directly comparable across models. The count importances scores can differ in scale between models due to differences in read depth between cell states. For visualization, for each cell state, we scale the count contribution score based on the percentile of per base scores (typically 0.1 for lower limit and 99.95 for upper limit). Values outside these bounds are clipped.

TF-MoDISco v0.5.14.0 (https://github.com/kundajelab/tfmodisco/tree/v0.5.14.0) (Shrikumar *et al.*, 2018) is used for *de novo* motif discovery i.e. to derive predictive consolidated motifs over a user-defined library of sequences and their associated contribution score profiles derived from a ChromBPNet model.  Briefly, TF-MoDISco first identifies contiguous subsequences with significant contribution scores in all the sequences scored through a modelFor each cell-state-specific model, the DeepSHAP scores (separately for count and profile) of sequences within peaks are cropped to 500 bp around the peak summit. TF-MoDISco is run with a maximum of 50,000 seqlets.

#### 2.8. Motif Analysis

##### 2.8.1. Motif Curation

After running TF-MoDISco on the first fold (fold 0) of each cell-state-specific model, we consolidated the motifs to obtain a set of predictive motifs across the entire reprogramming time course. We extracted the PFMs of TF-MoDISco motifs obtained from counts importance tracks across all models and clustered them using GimmeMotifs v0.15.3 (Bruse and van Heeringen, 2018) package's `gimme cluster` command with threshold 0.999. We then matched the clustered motifs using TomTom v5.3.0 (Gupta *et al.*, 2007) against a curated motif database (Vierstra *et al.*, 2020) consisting of motifs from JASPAR, HOCOMOCO, and SELEX. We used the arguments -no-ssc -oc . -verbosity 1 -text -min-overlap 5 -mi 1 -dist pearson -evalue -thresh 10.0 for TomTom. Following this, we excluded TF-MoDISco motifs with poor matches to existing motifs, and resolved ambiguities, in order to get a final set of 30 non-redundant predictive motifs whose presence is predicted to increase chromatin accessibility. For each motif, we chose as a representative the version from the cell type with the most instances. We then matched all MoDISco motifs against this set of 30 motifs and manually resolved minor consistencies and labeled examples which contained dimers of these 30 motifs.

##### 2.8.2. Predicted marginal ATAC-seq footprints associated with motifs

For each motif, we first selected its representative version (as defined in the previous section) corresponding to the cell type with the most instances. From this, we split the instances into 3 bins based on their log-odds score (low: 0.1-0.4, medium: 0.4-0.7, high: 0.7-1). For each cell state, we introduce these seqlets in the middle of random background regions for that cell state. We then compute the average predicted profile probability distribution over these sequences with and without Tn5 bias for each cell type using the trained bias and accessibility models.

##### 2.8.3. Read aggregation footprinting

To perform footprinting in a model-free manner, we used TF-MoDISco motifs for KLF and OCT-SOX from the xOSK state (C8), and AP1 motif from the fibroblast state (C1). We performed PWM scanning within peaks of each cell state. The motif instances were split into bins based on log-odds score thresholds. For each cell state and log-odds score bin, we aggregated the Tn5 insertions accounting for motif orientation. Specifically for each motif instance, we extracted 2000 bp centered at the motif, and normalized the total insertions to sum to 1 per locus. We then took the mean per position over all motif instances to obtain the final footprint for the cell state and log-odds bin combination, and plotted the central 100 bp.

##### 2.8.4. Motif Scanning

We extracted motif PWM scans for these motifs that lie within peaks by subsetting the pre-computed motif scanning track generated by Jeff Vierstra (Vierstra *et al.*, 2020). We further filtered PWM motif scan hits based on counts contribution scores, setting an empirical threshold for each motif as the 99th percentile of absolute summed contribution score of random regions within peaks with the same width as the motif, while taking into account the motif silhouette (defined as the maximum per position information content of the motif). We generated a matrix of peaks by motifs which was used for downstream analyses. We ran ChromVAR with default parameters on the binarized peaks x cells matrix to obtain a motif x cell matrix with ChromVAR motif deviation scores for each cell for each TF.

##### 2.8.5. Peak set x motif matrix

We summarized the peak by motif matrix into a count matrix of peak set by motif, allowing multiple hits for a given motif per peak. We first row normalized (over peak sets) this matrix, and set any element < 0.01 to 0. We then column normalized (over motifs) this matrix between 0 and 1. We then plotted this normalized peak set by motif matrix, plotting those entries with normalized scores greater than the first quartile of all non-zero scores.

##### 2.8.6. TF ChromVAR and expression pseudotime plots

We picked cells in cell states along the primary reprogramming trajectory (states 1,8,11-15), and ordered them by pseudotime computed using state 8 as root. We manually set fibroblasts (state 1) to have 0 pseudotime and iPSC (state 15) to have pseudotime 1, as they are disjoint from the other cells. The scATAC cells were matched to scRNA cells as those nearest in the Harmony corrected CCA space, with the constraint that both cells share the same cluster label. We then plotted heatmaps of ChromVAR score of the motifs arranged in increasing order of pseudotime. We also matched each motif with the TF from its motif archetype whose expression dynamics correlated with the ChromVAR score along pseudotime. To do this, we used the motif archetypes generated by Jeff Vierstra (Vierstra *et al.*, 2020). For smoothing, we computed a rolling average over every 250 cells, and also subsampled every 50 cells for plotting. For a few selected TFs, we plotted the ChromVAR score, gene expression and ArchR derived gene activity score along pseudotime. For smoothing, we twice computed the rolling average over 2000 cells. All values were min-max normalized to 0-1 for each TF and each modality (i.e. ChromVAR, expression, gene score).

##### 2.8.7. Sub-motif comparison

To compare the OCT-SOX motif instances between the xOSK state and iPSCs, we inspected the TF-MoDISco object run for each of the cell states. TF-MoDISco returns a list of motifs and the underlying instances (i.e. individual motif occurrences) that make up the motif. In addition, the instances are further clustered into sub-motifs. We focused on the OCT-SOX motif from both the TF-MoDISco objects. We excluded a sub-motif that corresponded to the OCT-only motif from the iPSC object. In order to compute a log-odds score for individual instances from both the sets, we used a log-odds matrix derived from the canonical OCT-SOX PFM from

(Avsec *et al.*, 2021), and used a uniform background of 0.25 for each base. For plotting, we used a minimum threshold of 4 for the log-odds scores. The results are unaffected by choice of threshold.

##### 2.8.8. OSK and AP1 motif affinity across peak sets

From the BPNet paper (Avsec *et al.*, 2021), we obtained the canonical ChIP-seq OCT-SOX, SOX and KLF motif PWMs. Separately, we obtained the AP-1 PWM from HOMER (GSE21512) (Heinz *et al.*, 2010). We found an appropriate log-odds threshold by scoring OCT-SOX, SOX and KLF seqlets in the xOSK state retrieved by TF-MoDISco and used the 10th percentile of scores as the threshold. This is required in order to focus on low affinity sites. We followed a similar approach for AP-1, but instead looked at instances in the Fibroblast cell state (which has high expression of the AP-1 factors). With these thresholds, we performed PWM scanning within each of the Early Transient and E-ON peak sets that are also xOSK peaks.

To refine our PWM hits using contribution scores, for each PWM hit of each TF, we computed the sum of xOSK counts contribution scores across the width of the motif, ignoring negative importance values. We then sampled 100,000 random interpreted regions of the same width and computed the sum of non-negative values for those, thereby obtaining a null distribution of these scores. We retained PWM hits whose contribution score was greater than the 99th percentile of the null distribution of scores. Finally, we removed instances of SOX and AP-1 motifs that overlap with OCT-SOX motif instances.

##### 2.8.9. Pseudo-binding saturation curves

The aim of this analysis is to study accessibility at sites activated by direct action of OSK at day 2 as a function of motif affinity and TF concentration. We focused on the OCT-SOX and

KLF motifs since they were the most prevalent in newly opened sites in the xOSK state. Since the xOSK state consisted of cells from both days 2 and 4, we first subsetted cells in the xOSK state to cells from day 2 only and ran the bulk ENCODE ATAC-seq pipeline v1.10.0 to call peaks. We restricted our analysis to xOSK peaks that were also (naive overlap) peaks in xOSK cells from day 2 only, and removed peaks that were open in fibroblasts. This peak set represents the first direct targets of OSK overexpression. We stratified the OCT-SOX motif instances in these peaks into low, medium and high affinity using the 40th and 90th percentiles of log-odds scores as breakpoints. We assigned each peak to a bin (low, medium, high) based on the affinity of the strongest motif present in it. In order to focus solely on the effect of the OCT-SOX heterodimer, we removed bins which contained a KLF motif. This resulted in a peak x OCT-SOX_affinity binary membership matrix with 3 columns: low, medium and high. We ran ChromVAR for each of these affinity bin associated peak sets separately. This returns a "motif activity" score for each cell for each affinity bin. For each affinity bin, we averaged ChromVAR deviation scores for each cell state along the primary reprogramming trajectory and min-max normalized the averaged scores. We repeated this analysis separately with the KLF motif wherein we removed peaks with the OCT-SOX motif.

##### 2.8.10. Per Gene Dynamic TF-Enhancer Network

For each gene of interest, we create a series of networks, one for each cell state, which consists of active enhancers putatively regulating the gene and transcription factors binding those enhancers. We consider all enhancers whose accessibility is correlated with the expression of the gene (absolute correlation>0.45 and FDR<1e-4.). Within a cell-state specific network, the active enhancer nodes correspond to the subset of all correlated enhancers that are accessible in that cell state. Edges are added between enhancer and TF nodes if a motif of that TF is present in the enhancer after thresholding by contribution scores (see Motif Curation).

In addition, we performed manual interventions to clean up the networks. First, we sub-selected a set of pertinent motif families. We removed SOX motifs that overlapped with OCT-SOX motifs, TFAP2 motifs that overlapped with KLF motifs, and KLF motifs that overlapped with CTCF motifs. Additionally, we marked as inactive motifs not recovered by TF-MoDISco in that cell state.

#### 2.9. Integrative Analysis

##### 2.9.1. Overlap with DHS Index

We downloaded the index of DNaseI Hypersensitive Sites (file DHS_Index_and_Vocabulary_hg38_WM20190703.txt.gz) from https://zenodo.org/record/3752359#.YcXz9y8RqTc (Meuleman *et al.*, 2020). For each of our 500 bp peaks, we considered it as present in the DHS Index if at least 100 bp was covered by the union of DHS Index regions using bedtools coverage.

##### 2.9.2. Bulk TF and histone ChIP-seq of human reprogramming

OSKM Day 2 bulk ChIP-seq data was taken from GSE36570 (Soufi *et al.*, 2012) and processed using the ENCODE ChIP-seq pipeline v1.3.6 with “pipeline_type” as “tf” and “peak_caller” as “macs2”. Reprogramming histone ChIP-seq data was taken from GSE62777 (Cacchiarelli *et al.*, 2015) and processed using the ENCODE ChIP-seq pipeline v1.3.6 with “pipeline_type” as “histone”. The signal fold-change tracks were used for visualization.

##### 2.9.3. MicroC Data

We downloaded Human Fetal Fibroblast (4DNFIPC7P27B) and H1-hESC MicroC (4DNFI2TK7L2F) data (Krietenstein *et al.*, 2020) from the 4D Nucleome portal. We visualized the MicroC data using BentoBox's (Kramer *et al.*, 2022) bb_plotHicSquare function with resolution=5000, norm="None" and zrange set to 5th and 95th quantiles of the non-zero counts for the NANOG locus.

#### 2.10. Single-nucleus Multiome processing

Processing for Multiome samples from days 1 and 2.

##### 2.10.1. Mapping

*ATAC*

ATAC processing was performed identical to scATAC-seq processing. We used the 10x Multiome barcode whitelist (737-arc-v1.txt.gz, copy available https://github.com/nanoporetech/sockeye/blob/5abeb52/data/737K-arc-v1.txt.gz) after reverse-complementing each barcode and appending the fixed 8bp prefix "CAGACGCG".

*RNA*

We used 10x Genomics Cell Ranger v6.0.2 (Zheng *et al.*, 2017) for read mapping and quantification to obtain the counts matrix. We used the GRCh38 2020-A reference. For each sample, we ran Cell Ranger count with --expect-cells=10000 --chemistry=ARC-v1 --include-introns. After processing each sample separately, we aggregated across samples using the Cell Ranger aggr command with --normalize=none. This ensures no reads are dropped by subsampling when aggregating counts across samples. The filtered_feature_bc_matrix is used for all downstream analyses.

##### 2.10.2. Quality Control

*ATAC*

We used ArchR v1.0.1 (Granja *et al.*, 2021) for initial quality control. We used the "createArrowFiles" function which takes as input all the fragment files from the mapping stage. In addition, we run the "addDoubletScores" function with parameters "k=10, knnMethod='UMAP', LSIMethod=1". These methods return quality control measures for each barcode as well as doublet enrichment scores. We use thresholds: total fragments >= 1000 and TSS enrichment >= 6. To remove doublets, we ran the AMULET method v1.1 (Thibodeau *et al.*, 2021) (https://github.com/UcarLab/AMULET/releases/download/v1.1/AMULET-v1.1.zip) on each sample. In addition to doublets called by AMULET, we called additional doublets based on read depth normalized overlaps (Number.of.Overlaps/Number.of.Valid.Reads from the OverlapSummary.txt file returned by AMULET) by using a manual knee-point detection strategy.

*RNA*

The RNA quality control is similar to what was done for scRNA-seq. One major difference was that nuclei with >0.5% OSKM transcript fraction were removed (compared to a threshold of >50% for scRNA-seq). This is because single-nucleus RNA-seq is not expected to contain Sendai transcripts which are primarily cytosolic. DoubletFinder v2.0.3 was run identical to the scRNA-seq case.

Finally, barcodes were filtered to those that passed QC in both ATAC and RNA individually, which resulted in 3217 nuclei from day 1 and 4161 nuclei from day 2.

##### 2.10.3. Dimensionality reduction, UMAP and cluster transfer

We used SnapATAC (Fang *et al.*, 2021) for dimensionality reduction and constructing the UMAP. We add the day 2 scATAC-seq cells along with day 1 and day 2 multiome nuclei. We analyzed the binary counts matrix over tiled 5kb genomic bins. Bins overlapping with the hg38 blacklist (Amemiya *et al.*, 2019) were omitted. We removed the top 5% of bins with the most cells. We performed dimensionality reduction with diffusion maps (runDiffusionMaps method with num.eigs=50). To alleviate the batch effect between scATAC-seq and multiome snATAC-seq, we used the Harmony algorithm (Korsunsky *et al.*, 2019) (runHarmony with eigs.dim=1:10). We called the runKNN method with eigs.dim=1:10, k=15 in order to construct the kNN graph from the first 10 Harmony dimensions. The runViz method with method="umap", eigs.dim=1:10 was used to create the UMAP. We next transferred cluster labels from scATAC-seqs to scMultiome cells. We used the FindTransferAnchors in Seurat on the first 10 Harmony embeddings with reduction='cca' followed by TransferData with weight.reduction='cca'.

#### 2.11. Multiome analyses

##### 2.11.1. Fibroblast-specific gene set

To obtain a set of fibroblast-specific genes for downstream analyses on the sn-Multiome data, we called differential genes from the scRNA-seq data. We used the FindMarkers function (min.pct = 0.25) in Seurat and compared the fibroblast cluster with the iPSC cluster to get fibroblast-specific differential genes. The function returned 1,113 genes at an adjusted *p-value* threshold of 0.01. In downstream analyses, for each nucleus, we computed the mean z-score across all genes in this set, which is called the "fibroblast gene program score".

##### 2.11.2. AP1 sequestration/retention plots

In these scatter plots, we compare the expression of fibroblast-specific genes (as detailed in the section "Fibroblast-specific gene set") to the estimated displacement/retention of AP1 TFs. For a given nucleus, the "AP1 sequestration score" is estimated by the normalized ATAC-seq insertions that fall in newly opened peaks (i.e. open in xOSK but not Fibroblast) with predictive AP1 instances. The normalization is performed by dividing the number of insertions by the total insertions in all ATAC-seq peaks. Analogously, the "AP1 retention score" is estimated as the normalized ATAC-seq insertions in Fibroblast peaks with predictive AP1 instances.
